# Identifying Intermolecular Interactions in Single-Molecule Localization Microscopy

**DOI:** 10.1101/2024.05.10.593617

**Authors:** Xingchi Yan, Polly Y. Yu, Arvind Srinivasan, Sohaib Abdul Rehman, Maxim B. Prigozhin

## Abstract

Intermolecular interactions underlie all cellular functions, yet visualizing these interactions at the single-molecule level remains challenging. Single-molecule localization microscopy (SMLM) offers a potential solution. Given a nanoscale map of two putative interaction partners, it should be possible to assign molecules either to the class of coupled pairs or to the class of non-coupled bystanders. Here, we developed a probabilistic algorithm that allows accurate determination of both the absolute number and the proportion of molecules that form coupled pairs. The algorithm calculates interaction probabilities for all possible pairs of localized molecules, selects the most likely interaction set, and corrects for any spurious colocalizations. Benchmarking this approach across a set of simulated molecular localization maps with varying densities (up to ∼ 50 molecules µm^*−*2^) and localization precisions (5 to 50 nm) showed typical errors in the identification of correct pairs of only a few percent. At molecular densities of ∼ 5-10 molecules µm^*−*2^ and localization precisions of 20-30 nm, which are typical parameters for SMLM imaging, the recall was ∼ 90%. The algorithm was effective at differentiating between non-interacting and coupled molecules both in simulations and experiments. Finally, it correctly inferred the number of coupled pairs over time in a simulated reaction-diffusion system, enabling determination of the underlying rate constants. The proposed approach promises to enable direct visualization and quantification of intermolecular interactions using SMLM.

## Introduction

Biomolecular interactions are fundamental to cellular physiology, governing critical processes such as cell signaling, gene regulation, and enzymatic catalysis. Understanding the spatiotemporal distribution of these interactions within cells is key to elucidating cellular function. However, direct observation of biomolecular interactions is a major technical challenge because these interactions occur on nanometer length scales – far below the diffraction limit of conventional optical techniques. While Förster Resonance Energy Transfer (FRET) (1; 2) can detect molecular colocalization using changes in either the intensity of fluorescence spectra or the excited state lifetimes, this method remains constrained by diffraction-limited spatial resolution. Biochemical approaches like coimmunoprecipitation (3; 4) or proximity labeling (5; 6) are powerful in identifying interaction partners, but they forfeit spatial information altogether. Thus, a critical need and opportunity exist to develop techniques to directly map biomolecular interactions with nanoscale resolution.

Single-molecule localization microscopy (SMLM) (7; 8) offers a promising approach to directly visualize and quantify biomolecular interactions at nanometer resolution. SMLM can determine the locations of individual fluorescently labeled molecules with a precision of 20-30 nm, approaching the length scale of intermolecular interactions. A key advantage of SMLM as compared to FRET is the nanoscale spatial mapping of molecular positions provided by SMLM. If supplemented with quantitative data on molecular binding, these spatial maps would allow probing how intermolecular coupling propensities depend on the heterogeneous local environment within cells. For example, SMLM could elucidate how interaction probabilities vary with local protein densities or how binding equilibria depend on access to protein nanodomains (9; 10; 11; 12; 13). Furthermore, by tracking dynamic interactions triggered by stimuli over time, SMLM could map spatiotemporal changes in reaction rates. Thus, adding quantitative binding information to super-resolved spatial maps provided by SMLM would result in a powerful approach to elucidate biomolecular interactions at the nanoscale in the complex cellular milieu.

Various methods have been proposed to analyze the spatial relationships between two kinds of molecules. For example, precise intermolecular distance measurements based on iterative localization of nominally identical protein complexes are possible (14; 15; 16). Unfortunately, these approaches cannot assign protein binding states in a spatially heterogeneous ensemble. For such datasets, methods have been developed to measure spatial correlations between molecular maps. For example, correlation-based metrics from diffraction-limited microscopy – Pearson correlation (17), cross correlation (18), and Mander’s overlap coefficient (19) – have been modified for use in SMLM (20; 21). However, these correlation-based methods are typically not constrained by the underlying reaction stoichiometry. Other recent methods include the development of an optimal transport approach to measure the distance between two distributions in a pixelated image (22), tessellation-based analysis to access the spatial organization of molecules by Voronoï diagrams (23; 24), and spatial point process based on Ripley’s K vector (25) to quantify the relative signal overlap between the two color channels. Although these methods are powerful in providing a relative measure of colocalization, they do not provide an absolute number of pairwise interactions.

Our goal was to count bound molecular pairs in the cell. This capability remains relatively underexplored in SMLM because it is challenging: SMLM localizations have finite precision and multiple sets of interacting pairs may be plausible. Here, we integrated these factors into a probabilistic model with the goal of determining both the absolute number and the fraction of coupled molecular pairs from two-color SMLM datasets. Given the observed inter-fluorophore distance and the corresponding localization precisions, we calculated the likelihood that the fluorescent tags resided within the expected interaction range for a bound complex. We then determined the most probable set of bound pairs in the SMLM image by maximizing the total probability over all putative pairs. Importantly, our approach focused on pairwise interactions by allowing each molecule to couple with at most one partner from the other channel, respecting stoichiometric constraints. Finally, to exclude random colocalizations, we estimated the number of spuriously paired molecules using Monte Carlo simulations of non-interacting particles at the relevant density and subtracted these chance events.

Evaluation of the overall analysis pipeline on simulated datasets with varying densities and localization precisions demonstrated excellent performance. Across the range of conditions tested, the fraction of correctly identified interacting pairs was typically over 95%. At the molecular density of ∼ 5-10 molecules per square micron and localization precision of 20-30 nm – conditions commonly encountered in SMLM measurements – recall exceeded 90%. Notably, our approach reliably distinguished non-interacting and bound proteins in both simulations and SMLM experiments. Furthermore, it accurately deduced the changing numbers of protein complexes that formed over time in a simulated reaction-diffusion system that evolved towards equilibrium; this allowed us to extract the underlying binding rate constants. These results demonstrate the capability of our probabilistic framework to identify molecular interactions using super-resolution microscopy data. This strategy has the potential to significantly advance our understanding of protein coupling at the subcellular level.

## Computational Methods

### Physical model of molecular interactions captured by SMLM imaging

We consider a prototypical system of two membrane proteins A and B that can bind reversibly: *A* + *B* ⇌ *AB*. Molecules of *A* and *B* are labelled with fluorescent tags of orthogonal spectral identities. In an SMLM image, the complex *AB* appears as a colocalization event between two spectrally distinct fluorophores. However, this colocalization is imperfect. Partly, this is due to the physical separation between the fluorophores (*d*_true_) labeling the two proteins. More importantly, *d*_true_ is a random variable because proteins may populate an ensemble of conformational states, and because dyes are typically attached to their target proteins via flexible tethers of finite length that undergo thermal fluctuations (Figure 1(a)). Furthermore, the detected positions of each fluorophore are also random variables. In the shot-noise-limited regime, the precisions of these localizations scale as 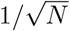 (26), where *N* is the number of detected photons. So, the observed distance *d*_obs_ is not only non-zero, but also is not necessarily equal to *d*_true_ (Figure 1(b)).

**Figure 1.**
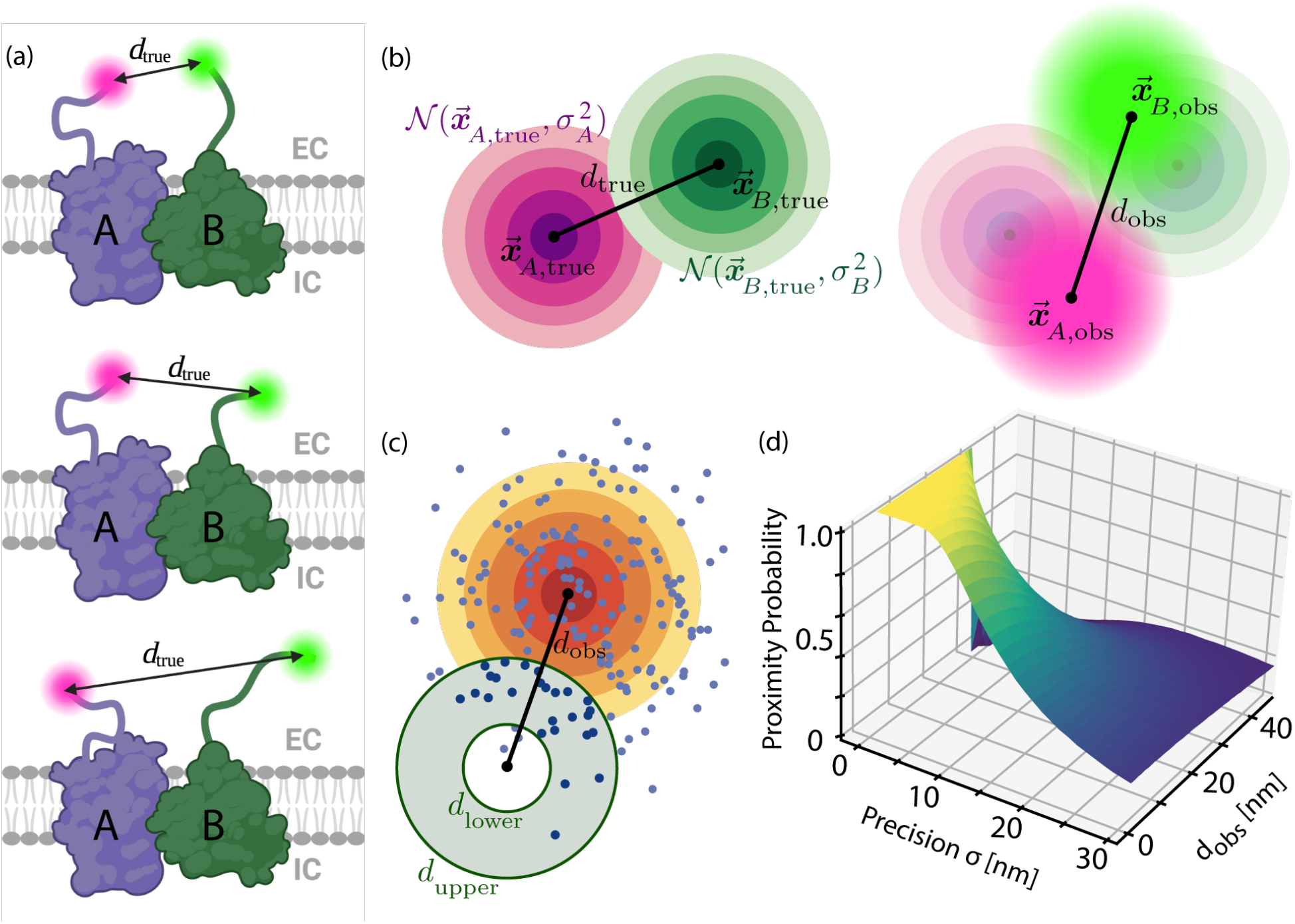
Defining and estimating the proximity probability *P*_prox_. (a) Illustration of three conformational states of a complex between two fluorescently labeled membrane proteins, *A* and *B*, with a variable distance *d*_true_ between the dyes, where the random variable *d*_true_ depends on the conformational state of the proteins and the flexibility of the dye linkers. EC and IC indicate extracellular and intracellular regions, respectively. (b) The observed distance *d*_obs_ is a random variable. Left: the magenta rings represent the Gaussian distribution centered at the true position 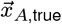 of the fluorophore *A* with localization precision σ_*A*_. Similarly, the green rings represent the Gaussian distribution at the location of the fluorophore *B*. The black line denotes the true distance *d*_true_. Right: the observed positions 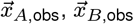 are random variables drawn from these distributions. Magenta and green halos illustrate signals obtained from the two fluorophores. The black line denotes the observed distance *d*_obs_. (c) Illustration of how the proximity probability *P*_prox_ is estimated by sampling points from a Gaussian distribution and counting the fraction of points that land in an annulus. The Gaussian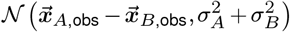, represented by the yellow rings, depends on the observed distance (black line) and localization precisions. The annulus, represented by the gray region, depends on a structural model of the complex *AB* and its fluorophores, where the lower bound *d*_lower_ and upper bound *d*_upper_ are constraints on *d*_obs_ consistent with colocalization. (d) *P*_prox_ can be estimated for arbitrary *d*_obs_, σ_*A*_, σ_*B*_, *d*_lower_, and *d*_upper_. For *d*_lower_ = 0, *P*_prox_ approaches 1 as *d*_obs_, σ_*A*_, σ_*B*_ → 0.

### Defining the probability that the observed distance is compatible with dimerization

We define the proximity probability as a metric of whether the positions and localization precision values of two emitters are compatible with a colocalization event. Assuming that the localized positions of the fluorophores labeling *A* and *B* follow Gaussian uncertainties σ_*A*_ and σ_*B*_, and are observed to be *d*_obs_ apart (Figure 1(b)), it can be shown (Supplementary Note 1) that the normalized squared observed distance follows a non-central chi-square distribution:

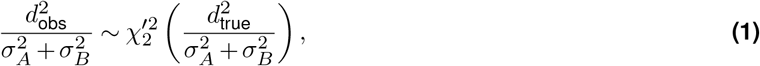

which fully describes ℙ (*d*_obs_ *d*_true_, *σ*_*A*_, *σ*_*B*_). However, we are interested in the inverse problem: given *d*_obs_, *σ*_*A*_, and *σ*_*B*_, can we infer whether *d*_true_ lies within a range given by a physical model? Rather than determining *d*_true_ from iterative SMLM measurements as done previously (14; 15; 16), we wish to estimate the proximity probability

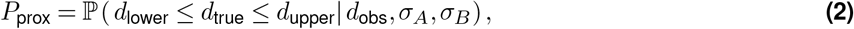

where *d*_lower_ and *d*_upper_ are constraints imposed by a structural model of the macromolecular complex *AB* and its fluorphores. Given two observed localizations, *P*_prox_ is the probability that the true distance *d*_true_ between the fluorophores lies within [*d*_lower_, *d*_true_]. When *d*_true_ lies within [*d*_lower_, *d*_true_], this proximity can originate from two scenarios: (1) molecular coupling, when the proteins A and B are bound in a complex *AB*, or (2) transient background pairing, when the uncoupled *A* and *B* diffuse close to each other by chance (Figure S3). We call the former “coupling”, and the latter “background pairing”. We refer to both cases jointly as “pairings” since they are indistinguishable at the level of individual colocalization events.

### Monte Carlo estimation of the proximity probability

We approximate *P*_prox_ by Monte Carlo sampling (Supplementary Note 2). Given observed positions 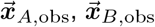 and localization precisions *σ*_*A*_, *σ*_*B*_, each Monte Carlo trial draws *N* points from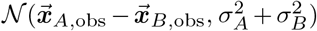. The fraction of points landing in the annulus with inner and outer radii *d*_lower_ and *d*_upper_ approximates *P*_prox_ (Figure 1(c)), which is highest when *σ*_*A*_, *σ*_*B*_ are small and the centroid of the Gaussian lies within [*d*_lower_, *d*_dupper_]. Figure 1(d) shows *P*_prox_ for a range of values of *d*_obs_ and *σ*_*A*_ = *σ*_*B*_, with *d*_lower_ = 0 and *d*_upper_ = 25 nm. Consistent with the ideal scenario of point particles localized with infinite precision, *P*_prox_ →1 as *d*_obs_ →0 and *σ*_*A*_, *σ*_*B*_ →0. In summary, our proposed Monte Carlo method estimates the probability that *d*_true_ lies within a range that is consistent with *AB* complex formation.

### Identifying pairings through Graph Matching Optimization (GMO)

Once a proximity probability is assigned to each pair of *A* and *B* using Eq. 2, we construct a bipartite graph that encodes all plausible coupling configurations. We select the most probable configuration by Graph Matching Optimization (GMO), the main idea of which is to represent a SMLM dataset as a bipartite graph with two sets of nodes, where each node in the set *V*_*A*_ represents a localization in the channel for *A* and each node in *V*_*B*_ represents a localization for *B*. We connect the nodes *A*_*i*_ and *B*_*j*_ with an edge if their proximity probability *p*_*ij*_ is nonzero, and assign *p*_*ij*_ as its edge weight. The most probable configuration is given by a selection of edges that maximizes the sum of *p*_*ij*_.

To save computational time, we only compute *p*_*ij*_ when 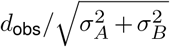 is less than a data-driven threshold *C* (Supplementary Note 3). Because the distribution in Eq. 1 has a thin tail, for a given distribution of localization precision values, *C* can be selected such that 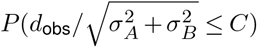 is arbitrarily close to one. For example, with 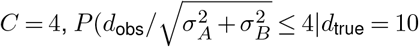, *σ*_*A*_ = *σ*_*B*_ = 15) = 99.96%, accounting for most relevant interactions (Figure S1(a)).

In summary, from a two-channel SMLM dataset (Figure 2(a)), we construct a weighted bipartite graph with two sets of vertices *V*_*A*_ and *V*_*B*_, and the set of all plausible edges *E* based on *P*_prox_ (Figure 2(b)). Each edge *e* is weighted by the proximity probability *w*(*e*) = *p*_*ij*_ of the two localizations.

**Figure 2.**
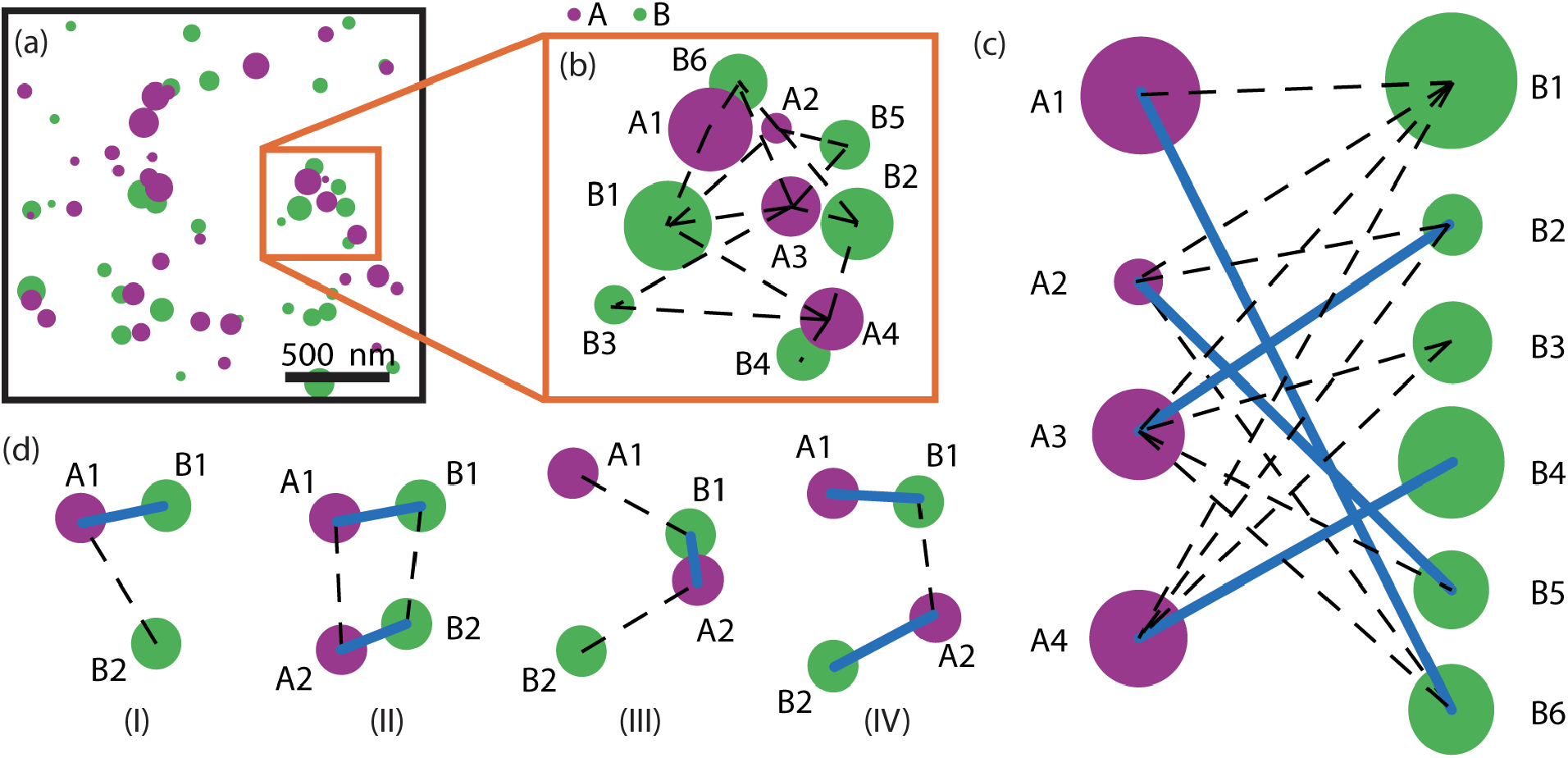
Graph Matching Optimization (GMO) selects the most probable configuration of molecular pairing. (a) A simulated SMLM image of proteins *A* (magenta) and *B* (green). Size of marker correlates with localization precision. (b) A connected component of the bipartite graph constructed for the dataset in (a). A node represents a localization of either *A* or *B*. An edge (dashed lines) connects *A*_*i*_ and *B*_*j*_ if their proximity probability *p*_*ij*_ is positive and their normalized observed distance is less than a data-driven threshold. (c) GMO selects a maximal weight matching (indicated by blue lines), a subgraph that maximizes the sum of *p*_*ij*_, and where each node is selected at most once. The matching represents a possible configuration of molecular pairing. (d) Matchings selected in four example scenarios. I: (*A*_1_, *B*_1_) is chosen over *A*_1_ and *B*_2_ if p_11_ > *p*_12_; II: (*A*_1_, *B*_1_) and (*A*_2_, *B*_2_) are pairs if *p*_11_ +*p*_22_ > *p*_12_ +*p*_21_; III: (*A*_2_, *B*_1_) is a pair if *p*_21_ > *p*_11_ +*p*_22_; IV: (*A*_1_, *B*_1_) and (*A*_2_, *B*_2_) are pairs if *p*_11_ +*p*_22_ > *p*_21_.

A possible configuration is represented by a *matching M* in the graph (*V*_*A*_, *V*_*B*_, *E, w*). A matching is a vertex-disjoint subset of edges; put simply, a matching pairs nodes such that each node is used at most once. In our context, (*A*_*i*_, *B*_*j*_) is selected for the matching *M* if and only if the localizations *A*_*i*_ and *B*_*j*_ are paired. For example, in the matching shown in Figure 2(c), the (blue) edges represent pairs. We emphasize that not every pairing is necessarily a molecular coupling event as the molecules could be near each other by random chance. We account for this effect in the next section.

The most probable configuration is given by a matching that maximizes the total sum of proximity probabilities. Thus, the objective is to find a set of edges *M*^∗^ that maximizes the sum of edge weights *p*_*ij*_, subject to the constraint that each node is selected at most once. This can be achieved by the following combinatorial optimization problem (27):

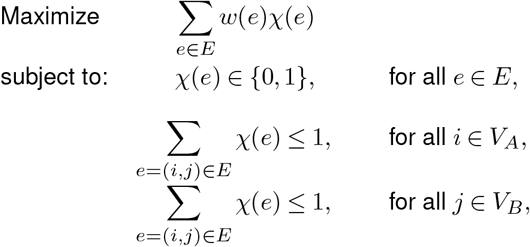

where each node *i* ∈ *V*_*A*_ represents a molecule of *A, j* ∈ *V*_*B*_ represents a molecule of *B*, and an edge *e* may connect the nodes *i* and *j*, with χ (*e*) = 1 indicating that this edge is selected for the matching *M*^∗^. The last two constraints ensure that each node is selected by at most one edge. Figure 2(d) shows several simple examples of this optimization step. In all cases, the selected matchings (blue edges) maximize the sums of proximity probabilities *p*_*ij*_.

### Iterative Monte Carlo Estimation of Molecular Couplings and Background Pairings (iMEC)

The matching obtained by GMO represents the most probable configuration of molecular pairings, which consists of both bona fide couplings and pairings by random chance, i.e., “background pairings” (Figure S3). We estimate the number of background pairs, and thus the number of coupling events, by iterative estimation. We call this process Iterative Monte Carlo Estimation of Molecular Couplings and Background Pairings (iMEC).

To illustrate the method, consider the extreme scenario where *A* and *B* do not interact at all (*K*_binding_ = 0) but the densities of both species are sufficiently high. If all localizations from *A* and *B* are uniformly distributed, GMO would return a non-empty matching. We would interpret all these edges as background pairs.

More generally, consider a SMLM dataset with *N*_*A*_ localizations in the spectral channel for species *A* and *N*_*B*_ localizations in the spectral channel for species *B*, where GMO returns a maximal weight matching *M*^∗^ with 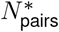 edges. We assume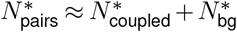, where 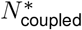 is the number of true molecular coupling events and 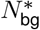 is the number of background pairings. If 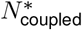 is known, we can estimate 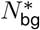 by simulating a spatial Poisson process using 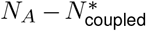 copies of A and 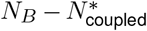 copies of *B* and applying GMO. However, to infer the unknown quantity 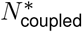, we estimate 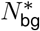 by an iterative procedure (Figure 3(a)). By estimating 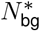, we can infer 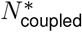.

**Figure 3.**
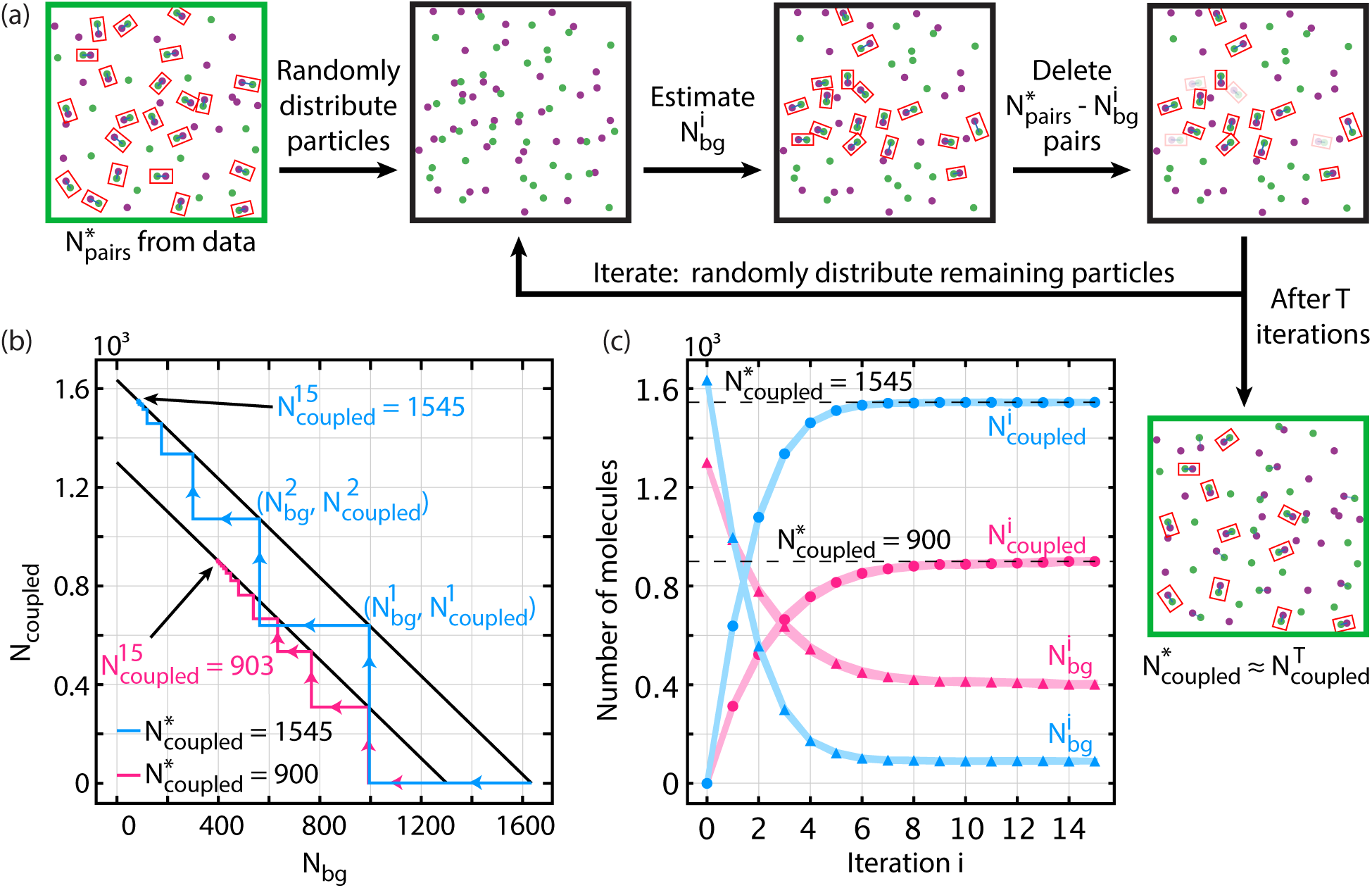
Iterative Monte Carlo Estimation of Molecular Couplings and Background Pairings (iMEC) estimates the number of background pairs among a configuration of molecular pairing. (a) Process overview of iMEC: starting with the most probable configuration with 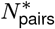 pairings from GMO, distribute the localizations in an experimentally imaged region uniformly random to estimate the number of background pairs 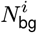. For the next iteration, repeat the process with 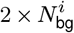 fewer localizations, which is equivalent to deleting 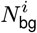 pairs from the dataset. The number of couplings 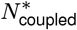 is estimated successively by 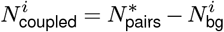. (b) Two example progressions shown on a plot of 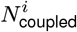 versus 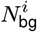. Of the 2000 localizations from *A* and 2000 localizations from *B*, the number of true molecular couplings were 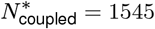 (blue) and 900 (pink). The lines of slope *−*1 denote the conserved sum 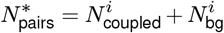. Arrows show iterative progression to convergence. After 15 iterations, the estimates were 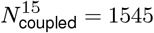 (blue) and 903 (pink). (c) 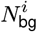 and 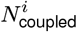 for the examples shown in (b), averaged over 10 Monte Carlo trials.

For the first iteration of this iterative procedure, we assume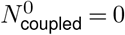, and simulate a spatial Poisson process with all *N*_*A*_ localizations of *A* and *N*_*B*_ localizations of *B*. GMO provides an initial estimate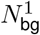, from which the number of putative true couplings can be inferred as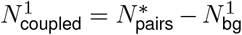. If 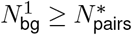, the number of pairs can be explained by chance alone. Otherwise, if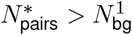, then there are at least 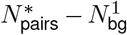 real coupling events. In the second iteration, we exclude the putative true couplings by applying GMO to a spatial Poisson process with 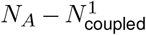 and 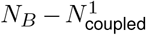 localizations to re-estimate the number of background pairs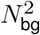. We then iterate this process, each time reducing the pool of potential background pairings by the number of couplings inferred in the previous interation round (Figures 3(b) and 3(c)). The process can be stopped after it meets a convergence criterion, or, in our case, after a fixed number of iterations. Details and pseudo-code are available in Supplementary Note 4.

Figure 3(c) shows 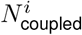 and 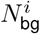 converging to 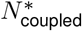 and 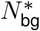, respectively, after 7 iterations. If 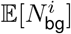 is non-increasing, the algorithm converges. Indeed, in all simulations, convergences similar to those shown in Figure 3(c) were observed.

## Results

The algorithm described above was evaluated across a range of possible scenarios. First, the performance was benchmarked via Monte Carlo simulations across varying experimental conditions – localization precision and density values. Next, the method was validated using experimental and simulated data of both non-interacting and interacting particles at equilibrium. Finally, we tested our approach on simulated non-equilibrium reaction-diffusion dynamics with the goal of capturing temporal changes in molecular couplings.

### Algorithm performance across varying localization precision and density

We evaluated the algorithm’s performance over a range of localization precision values, *σ*_*i*_, and molecular densities, *ρ*, using simulated data with equal numbers of species *A, B*, and *AB*. Two metrics were assessed: recall rate, measuring the fraction of true positives detected, and error rate, quantifying accuracy in estimating the number of true couplings. These two metrics evaluate the two facets of the algorithm: identifying likely couplings through GMO (recall), and inferring the number of true couplings via iMEC (error). The Methods section contains a description of how we generated the simulated datasets; full implementation details and parameters can be found in Supplementary Notes 6 and 7.

At the density of ρ = 5 molecules/µm^2^, recall rates remained above 80% even when localization precision reached up to 50 nm (Figure 4(a)). At 20 molecules/µm^2^, recall was above 80% for localization precision values below 25 nm (Figure 4(a)). We also did an analogous analysis across densities (Figure 4(b)). At the localization precision *σ*_*A*_ = *σ*_*B*_ = 10 nm, recall was greater than 90% for densities up to 50 molecules/µm^2^. Retaining recall greater than 80% at *σ*_*A*_ = *σ*_*B*_ = 30 nm required molecular densities below 15 molecules/µm^2^. These results demonstrate robust detection of molecular pairs with GMO at typical SMLM imaging conditions.

**Figure 4.**
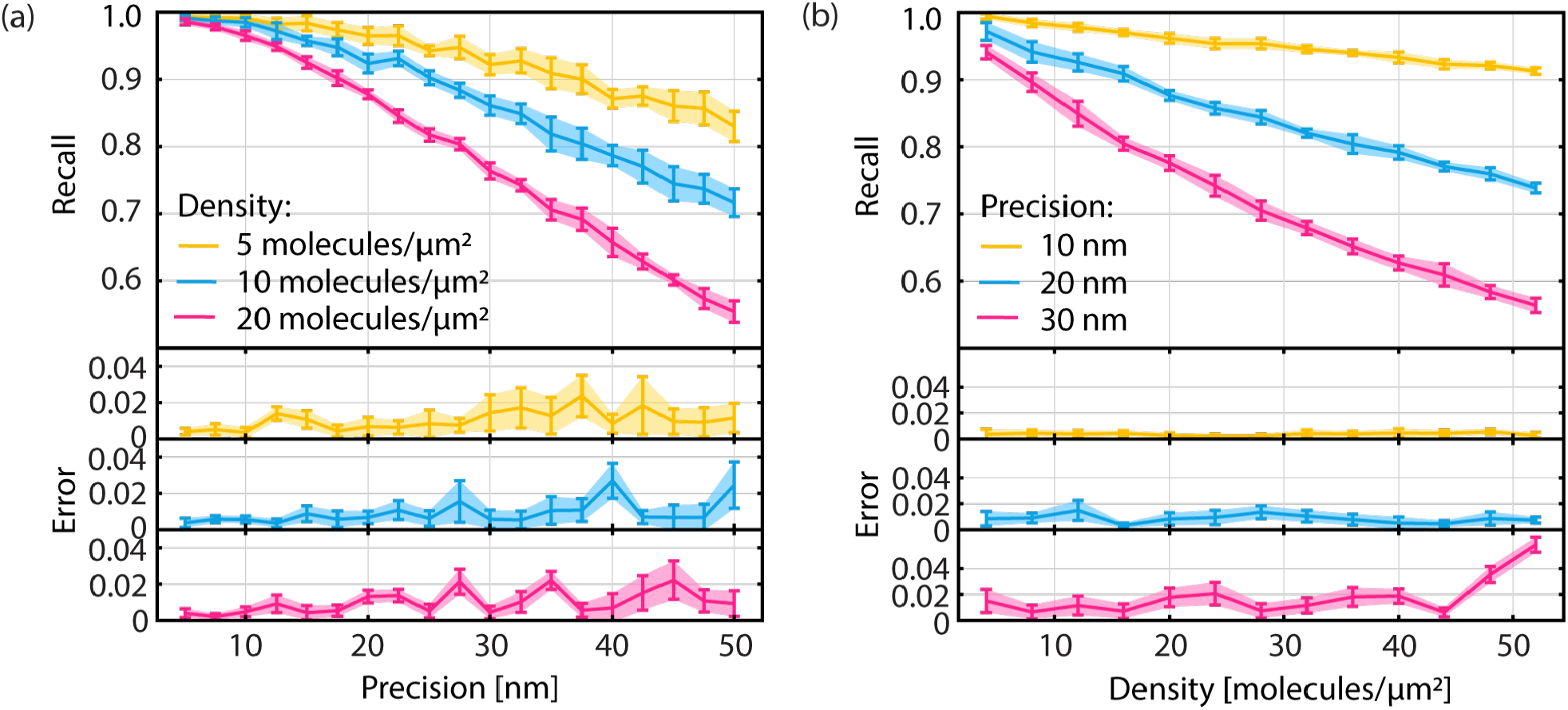
Algorithm performance on simulated datasets. (a) Recall rate and error rate as a function of localization precision. Precise localizations lead to better performance. (b) Recall rate and error rate as a function of molecular density. Recall is better at lower densities, while error remains mostly unchanged. Error bars indicate *±*1 standard deviation across 10 trials of iMEC.

Estimation errors were less than 5% across all conditions (Figure 4), with minimal dependence on density from 4-50 molecules/µm^2^ (Figure 4(b)). Errors increased slightly at lower localization precision but remained below 4% even at a marginal localization precision of 50 nm (Figure 4(a)). These results demonstrate reliable estimation of the number of true pairs by iMEC. We also found that iMEC consistently outperformed the naive approach of selecting colocalizations based on minimal distances; see Supplementary Notes 8 for details.

In summary, the algorithm achieved accurate detection of molecular couplings under typical SMLM experimental conditions (localization precision 5-50 nm and densities ≤ 50 molecules/µm^2^), which establishes its suitability for application to experimental SMLM datasets. As expected, recall decreased as localization precision deteriorated and as density increased.

### Algorithm validation at equilibrium using simulations and experiments

We validated the algorithm by asking whether it could accurately quantify the fraction of molecular couplings at equilibrium. Experimental SMLM data were acquired using two protein populations. The first population consisted of HaloTag linked to the N terminus of a β2 adrenergic receptor (Halo-β2AR) and SNAP-tag linked to a CaaX box sequence (SNAP-CaaX) that attaches to the plasma membrane. This population acted as a non-interacting negative control. The second protein population consists of a transmembrane helix with HaloTag on the N terminus and SNAP-tag on the C terminus (Halo-TM-SNAP), which acts as a positively interacting control (Figure 5(a)). We hypothesized that the Halo-TM-SNAP data would show significant colocalization between the spectral channels, while the Halo-β2AR/SNAP-CaaX pair would mimic randomly distributed non-interacting proteins. Confocal imaging confirmed expression and membrane localization of both constructs (Figure 5(b)). SMLM imaging (Figure 5(c)) and subsequent analysis by the GMO and iMEC pipeline showed significantly more coupled pairs for the positive control (21 ± 5%) compared to the negative control (6 ± 2%) (Figure 5(d)). These results demonstrate successful quantification of equilibrium interactions from experimental data. Details of experimental and imaging protocols can be found in Supplementary Note 9; parameters used for analysis are available in Supplementary Note 7.

**Figure 5.**
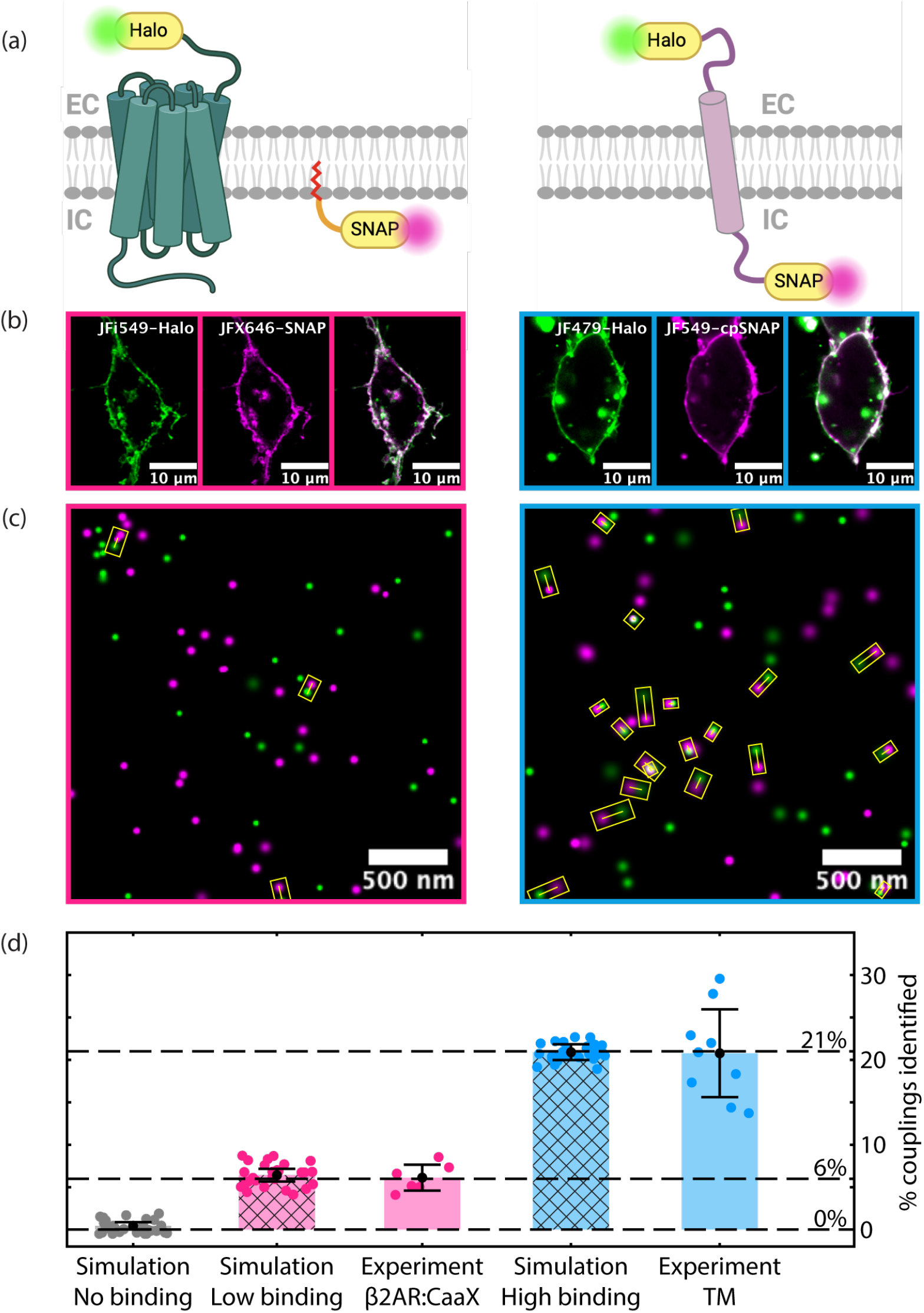
Validation of the algorithm on simulated and experimental SMLM data. (a) Schematics of Halo-β2AR/SNAP-CaaX (negative control) and Halo-TM-SNAP (positive control) – two test systems of membrane proteins. Halo-TM-SNAP was expected to show significantly more colocalization events than Halo-β2AR/SNAP-CaaX. (b) Confocal images of Halo-β2AR/SNAP-CaaX (left) and Halo-TM-SNAP (right) expressed in HEK 293FT cells. Confocal imaging demonstrates localization of constructs to the plasma membrane but does not provide information about molecular coupling. (c) Sample regions of experimental SMLM images for Halo-β2AR/SNAP-CaaX (left) and Halo-TM-SNAP (right). Paired molecules are highlighted with yellow rectangles. (d) Percentage of identified colocalizations for Halo-TM-SNAP, Halo-β2AR/SNAP-CaaX, and simulated datasets with different binding strengths. In the simulations, coupling percentages are set at 0% for no binding, 6% for Halo-β2AR/SNAP-CaaX (low-binding), and 21% for Halo-TM-SNAP (high-binding), the latter two derived from mean values calculated from experimental data. Each dot represents either a cell or a simulated dataset; bars represent averages over all cells/simulated datasets. Error bars indicate *±*1 standard deviation. Dashed horizontal lines denote the ground truth for no-binding, low-binding, and high-binding scenarios.

We also analyzed simulations of the reaction *A* + *B* ⇌ *AB* at equilibrium, where the number of couplings was dictated by the equilibrium constant K_eq_ (Supplementary Note 12). Our algorithm reliably recovered the true number of *AB* complexes across low ( ∼ 0.005) and high ( ∼ 0.07) *K*_eq_ values (Figure 5(d)). In addition, we investigated the algorithm’s performance across a broad range of densities (at different *K*_eq_ values) and found that the output of the algorithm matched the theoretical expected values (Figure S16). Finally, we evaluated and demonstrated the consistency of our algorithm across different density ratios between the two channels, a common challenge in colocalization analysis (17; 25; 24) (Supplementary Note 13). Taken together, these results validated our pipeline’s capability of accurately quantifying molecular couplings at equilibrium from both simulated and experimental SMLM data.

### Algorithm validation on non-equilibrium dynamics

Next, we tested whether our method could analyze data from cells in non-equilibrium states. Recent advances in time-resolved techniques, like time-resolved cryo-vitrification (28; 29; 30; 31), allow precise stimulation and fixation of cells at defined timepoints followed by super-resolution imaging (Figures 6(a) and 6(b)). Applying our pipeline to these static images can connect them into a dynamic sequence, revealing temporal information on molecular binding.

**Figure 6.**
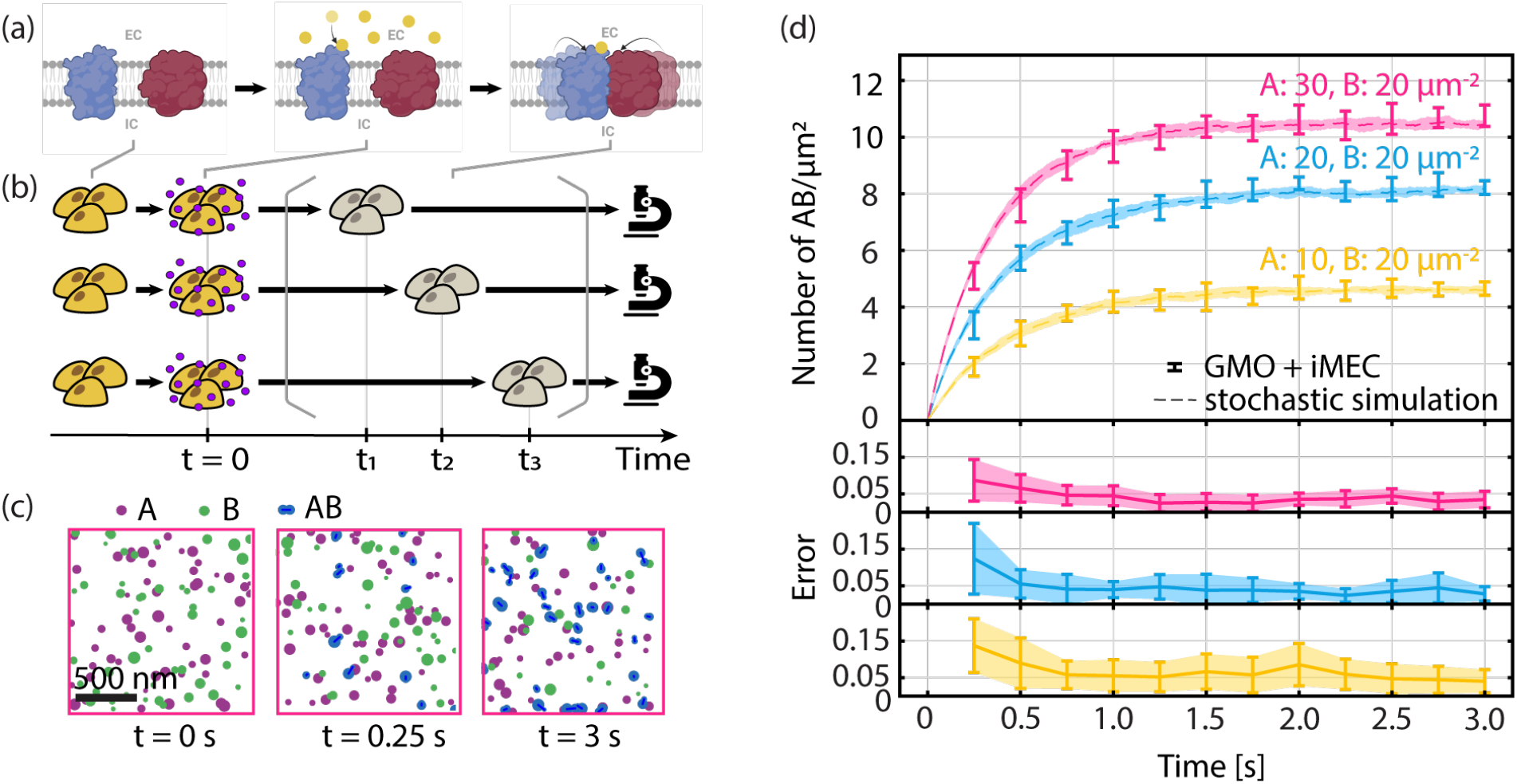
Validation of the algorithm on simulated non-equilibrium binding dynamics. (a) Schematic of a hypothetical biological system: following ligand stimulation at *t* = 0, membrane protein *A* (blue) becomes active and can bind to membrane protein *B* (red). EC and IC indicate extracellular and intracellular regions, respectively. (b) Schematic of a hypothetical experimental protocol: ensembles of cells are stimulated with ligand at *t* = 0 and are fixed at times *t*_1_, *t*_2_, *t*_3_ for SMLM imaging. (c) Snapshots from a simulation modeling protein binding, *A*+*B* ⇌ *AB. AB* shown indicate true binding. Snapshots were taken at *t* = 0, 0.25 s, and 3 s after ligand addition. (d) Top: Density of *AB* identified by the algorithm at various time points (each time point consists of 10 simulated datasets). The variations in the shaded regions originate from stochastic simulations, while the variations in the error bars are attributed to the inference algorithm. Bottom: Error rate at various time points. Error bars indicate *±*1 standard deviation.

To evaluate the method’s utility for reaction kinetics, we simulated a model of the reaction *A* + *B* ⇌ *AB* (details in Supplementary Note 6B), extracted the positions of molecules at select timepoints to generate simulated SMLM datasets (Figure 6(c)), and analyzed them using our algorithm. We accurately reproduced the number of complexes across timepoints and densities, with ∼ 10% average error (Figure 6(d)). The slightly higher error relative to those of Figure 4 likely stemmed from fewer total molecules in these simulations. Errors were also larger at earlier timepoints, again due to fewer complexes. In summary, our algorithm captured the dynamics in non-equilibrium simulations, demonstrating its potential for analyzing time-resolved SMLM measurements of cellular processes.

We also estimated the rate constants by fitting the density of coupled pairs to the concentration of *AB* over time (Supplementary Note 14). Our estimates of 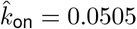 molecules^−1^µm^2^s^−1^ and 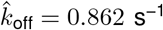 closely match the theoretical input values of *k*_on_ = 0.0549 molecules^−1^µm^2^s^−1^ and *k*_off_ = 1 s^−1^. This and the results shown in Figure 6(d) together demonstrate the method’s potential to quantitatively track reaction progression.

## Discussion and Conclusion

Super-resolution microscopy enables nanoscale mapping of cellular components but cannot directly discern functional binding states. We proposed a pipeline that bridges this gap by statistically deducing protein interactions from single-molecule localization data. Performance assessment showed strong recall and small error rates under typical SMLM conditions. The technique was also able to differentiate between positive and negative experimental controls – a result that was supported by Monte Carlo simulations. When applied to non-equilibrium reaction-diffusion simulations, the approach reliably recovered the temporal evolution of molecular coupling. Taken together, these results validate the algorithm’s capacity to infer bona fide couplings from SMLM data in both steady-state and transient cellular processes.

The key innovation of our approach is the ability to estimate the absolute number of molecular couplings. Our algorithm consists of three parts. First, we compute the proximity probabilities between pairs of molecules, indicating potential interactions. Second, GMO identifies the most probable configuration of pairings by maximizing the sum of proximity probabilities. Finally, iMEC estimates the subset of pairings arising from true interactions rather than random colocalization. Notable strengths of our approach include: (1) elucidation of position-dependent, molecular-scale interactions undetectable by diffraction-limited techniques; (2) the use of a biophysical probabilistic model to provide a robust foundation for statistical inference; (3) rigorous correction for chance colocalizations; and (4) demonstrated accuracy on equilibrium and kinetic binding data across a broad range of densities and localization precision values.

However, limitations also exist. While the algorithm can estimate the total number and percentage of true interactions, it cannot accurately determine whether a specific interacting pair represents a true coupling or a background pairing. In addition, the approach relies on the quality of the input SMLM data. Factors that limit SMLM performance, including drift, optical aberrations, incomplete labeling, premature photoactivation or photobleaching, also constrain the quality of algorithm’s output. Accurate detection of molecular coupling is particularly challenging because colocalization probability scales with the square of the fraction of detected molecules (Supplementary Note 11C). Moreover, complex spatial distributions of molecules inside cells typically deviate from a uniform distribution, which makes the Monte-Carlo-based inference prone to error. Addressing these limitations is an opportunity for future work. Additional future directions could explore algorithmic alternatives to GMO and iMEC, expand the approach to handle 3D SMLM images, analyze multi-channel SMLM datasets beyond two colors, and address heterogeneous and higher-order interactions.

In summary, the broadly applicable framework presented here infers protein binding from single-molecule localization data, enabling quantification of the spatial relationships of molecular coupling. This versatile new platform has the potential to elucidate the intricate protein interaction landscapes governing diverse cellular functions. We envision the algorithm finding biological applications in probing dynamic protein interactions underlying key cellular processes including transmembrane signaling, gene regulation, and enzymatic catalysis.

## Materials and Methods

### Implementation of GMO

In constructing the weighted bipartite graph for a dataset, we only considered pairs of localizations that are not too far apart. We first determined the threshold parameter based on the distribution of localization precision values (Supplementary Note 3). For pairs of localizations meeting this threshold, we estimated the proximity probability *p*_*ij*_ (Supplementary Note 2). Using these probabilities, the bipartite graph was built, and the edge weights were taken to be ⌊10^5^*p*_*ij*_⌋. The resulting graph was fed into NetworkX’s function `max_weight_matching` (32), which then returned a graph matching that maximized the sum of proximity probabilities.

### Implementation of iMEC

iMEC was implemented in Python. Detailed description can be found in the corresponding section in the main text and (Supplementary Note 4).

### Generating simulated datasets

To simulate a system at equilibrium, we instantiated molecules of *A, B*, and *AB* uniformly in space, and symmetrically split any *AB* into a localization for *A* and a localization for *B*, at a random distance drawn from Unif(0, *d*_true,max_), where *d*_true,max_ = 10 nm. See (Supplementary Note 6A) for implementation details. Localization precision values were either set at a fixed value, or drawn from the experimental localization precision distribution in Supplementary Note 6C with a cutoff at 40 nm. All parameters used are available in Supplementary Note 7.

### Generating simulated non-equilibrium binding dynamics

The non-equilibrium dynamics in Figure 6 were generated by a stochastic particle-based reaction-diffusion simulation, implemented using ReaDDY (33). Implementation details, including parameters, are available in (Supplementary Note 6B).

### Cell culture and transfection

Two cell lines were used: Human Embryonic Kidney 293FT (HEK293FT) cells and HEK293 cells with a CRISPR/Cas9 knockout of the G-protein Gs (HEK293 ΔGs). Cells were cultured in treated cell culture flasks with Dulbecco’s Modification of Eagle’s Medium (DMEM) with 4.5 g/L glucose, L-glutamine, & sodium pyruvate. Penicillin-Streptomycin solution was added to prevent bacterial contamination. Plasmid transfections were done in either 6-well or 12-well plates using either Lipofectomene 3000 or Polyplus JetOptimus according to manufacturer protocol. Details about the cell lines and protocols are available in (Supplementary Note 9A). A full list of plasmids used can be found in Supplementary Note 9A.

### Confocal imaging

Cells were seeded onto glass-bottom dishes and labeled with fluorescent dyes. Identities of dyes used in specific experiments can be found in relevant figures. For fluorescent labeling, a dye solution in cell media was prepared at a final concentration of 2 µM, and cells were labeled by incubation in dye solution for 15 minutes. Cells were subsequently washed to remove non-specifically bound dye. Dishes were imaged on a Zeiss LSM 880 confocal microscope using a 63x / 1.40 NA oil objective. Full labeling and imaging parameters are available in Supplementary Note 9B. Confocal images were adjusted for brightness and contrast using Fiji software.

### SMLM imaging

Cells were labeled with super-resolution compatible fluorescent dyes with a protocol modified from “Confocal Imaging.” After labeling and washing, cells were replated onto pre-cleaned glass coverslips. Once cells adhered, they were fixed with a solution of 4% PFA in PBS. Coverslips were mounted onto glass slides with a drop of Fluoromount-G as the mounting medium. Coverslips were sealed prior to imaging. Samples were imaged on a Zeiss Elyra microscope using a 63x/1.40 NA oil objective. Widefield image stacks were collected in two emission channels corresponding to red and far red dyes. Over the course of imaging, dyes were stochastically activated with a 405 nm laser. Details can be found in Supplementary Note 9C.

### Processing of SMLM images

Single-molecule localizations were processed in Zeiss Zen software. Localizations were grouped to aggregate localizations from a single dye molecule spread over multiple consecutive frames. They were then filtered, drift-corrected, and clusters were identified and subsequently removed using DBSCAN (34), implemented using the `sklearn.cluster.DBSCAN` package, with parameters `eps = 75` nm and `min_samples = 10`. Finally, the image was divided into smaller subregions, of which any dense subregions were excluded from analysis. See Supplementary Note 10A for details.

### Analyzing SMLM images

Each subregion was analyzed using the GMO+iMEC pipeline (Figure S4), and the percentage of couplings in the subregion was calculated using Eq. 5 below. Each dot in Figure 5(d) represents the average percentage of couplings for each cell. See details in Supplementary Note 10B.

### Evaluation metrics

Two metrics were used to evaluate the performances of our methods. *Recall* was used to measure the percentage of true couplings successfully retrieved (true positive or TP) relative to false negatives (FN) in GMO:

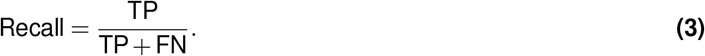

We used the *error rate* to measure the performance of iMEC:

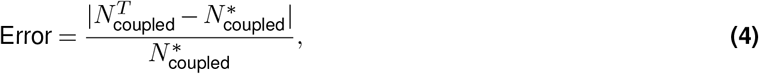

where 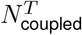 was the output from iMEC after *T* iterations, and 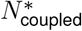 was the number of true couplings in the simulated dataset. To compare datasets with different densities (e.g., in Figure 5(d)), we defined for each region

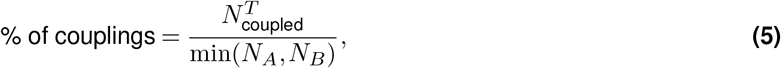

where *N*_*A*_ was the number of localizations corresponding to *A*, and *N*_*B*_ was the number of localizations corresponding to *B*.

## Acknowledgements

This research was supported by the Aramont Fellowship Fund for Emerging Science Research, NIH Research Project Grant R01 GM146791 and R21 GM146127 and startup funds from Harvard University. X.Y. and P.Y.Y. were supported by NSF-Simons Center for the Mathematical & Statistical Analysis of Biology (DMS-174269) and the Harvard Quantitative Biology Initiative. A.S. was supported in part by Harvard Qbio Student Award and Simmons Award from Harvard Center for Biological Imaging. The authors thank the Harvard Center for Biological Imaging (RRID:SCR_018673) for infrastructure and support. The computations in this paper were run on the FASRC Cannon cluster supported by the FAS Division of Science Research Computing Group at Harvard University. The authors thank Asuka Inoue at Tohoku University for providing the HEK293 ΔGs cells. The authors thank Johannes Broichhagen at Leibniz-Forchungsinstitut for Molecular Pharmocology for providing the Halo-TM-SNAP DNA construct. The authors thank Simon Merminod, Bridget Queenan, Samuel Kou, Jeremy Conway, Ami Thakrar, Daphne-Eleni Archonta and Jenny Hong for helpful discussions. We thank Rachelle Gaudet, Daniel Needleman and Douglas Richardson for feedback on the manuscript.

## Author Contributions

X.Y., and M.B.P. conceived the project; X.Y. developed the method and associated software; P.Y.Y. developed the stochastic simulation and associated software; A.S. performed the experiments and collected data; X.Y., P.Y.Y., A.S., and M.B.P. analyzed data; A.S. and S.A.R. contributed to single-molecule image processing and analysis. X.Y., P.Y.Y., A.S., and M.B.P. wrote the manuscript. M.B.P. supervised the research.

