## Supplementary Information for "Identifying Intermolecular Interactions in Single-Molecule Localization Microscopy"

### Supplementary Information Content

**1 Derivation of the distribution for the observed distance between two fluorophores** **2**

**2 Estimating the proximity probability of a pair** **3**

**3 Data-driven selection of a threshold parameter for $d_{\text{obs}}$** **4**

**4 Estimating the true number of molecular couplings and the number of pairings from transient proximity** **5**

**5 Pipeline to estimate the number of molecule couplings in an SMLM image** **7**

**6 Generating simulated SMLM datasets** **8**

**12 Validation at chemical equilibrium** **23**

**13 The algorithm can estimate the level of coupling accurately across different density ratios** **24**

**14 Inferring rate constants from kinetics data** **25**
- A Analytical solution . . . 25
- B Inferring rate constants . . . 25

### Supplementary Information

#### Supplementary Note 1: Derivation of the distribution for the observed distance between two fluorophores

Here we derive the distribution of the observed distance between two fluorophores in 2D SMLM. We show that the normalized observed distance squared and the normalized true distance squared follow a non-central chi-square relationship:

$$\frac{d^2_{\text{obs}}}{\sigma_A^2+\sigma_B^2} \sim \chi_2'^2\left(\frac{d^2_{\text{true}}}{\sigma_A^2+\sigma_B^2}\right). \tag{S1}$$

Consider two spectrally distinct fluorophores $A$ and $B$, whose true positions are $(A_x, A_y)$ and $(B_x, B_y)$ respectively. Assume their observed positions $O_A = (O_{Ax}, O_{Ay})$ and $O_B = (O_{Bx}, O_{By})$ are drawn from Gaussian distributions centered at their true positions. For simplicity, we assume that the localization uncertainties are radially symmetric. Let the localization precision values for $A$ and $B$ be respectively $\sigma_A$ and $\sigma_B$ (1).

From $O_A \sim \mathcal{N}((A_x, A_y), \sigma_A^2)$ and $O_B \sim \mathcal{N}((B_x, B_y), \sigma_B^2)$, we rescale to obtain

$$\frac{(O_{Ax}-O_{Bx})-(A_x-B_x)}{\sqrt{\sigma_A^2+\sigma_B^2}} \sim \mathcal{N}(0,1) \quad \text{and} \quad \frac{(O_{Ay}-O_{By})-(A_y-B_y)}{\sqrt{\sigma_A^2+\sigma_B^2}} \sim \mathcal{N}(0,1). \tag{S2}$$

Converting to chi-square distributions, we get

$$\frac{(O_{Ax}-O_{Bx})^2}{\sigma_A^2+\sigma_B^2} \sim \chi_1'^2\left(\frac{(A_x-B_x)^2}{\sigma_A^2+\sigma_B^2}\right) \quad \text{and} \quad \frac{(O_{Ay}-O_{By})^2}{\sigma_A^2+\sigma_B^2} \sim \chi_1'^2\left(\frac{(A_y-B_y)^2}{\sigma_A^2+\sigma_B^2}\right), \tag{S3}$$

where $\chi_k'^2(\lambda)$ is the non-central chi-square distribution with degree of freedom $k$ and noncentrality parameter $\lambda$. Note that the noncentrality parameter $\lambda_{\sigma_A,\sigma_B,d_{\text{true}}}$ depends on $\sigma_A$, $\sigma_B$ as well as $d_{\text{true}}$. Then the square of the observed distance, normalized by $\sigma_A^2+\sigma_B^2$, follows a non-central chi-square distribution with degree of freedom $k = 2$ (2):

$$\frac{d^2_{\text{obs}}}{\sigma_A^2+\sigma_B^2} = \frac{(O_{Ax}-O_{Bx})^2}{\sigma_A^2+\sigma_B^2} + \frac{(O_{Ay}-O_{By})^2}{\sigma_A^2+\sigma_B^2} \sim \chi_2'^2\left(\frac{(A_x-B_x)^2}{\sigma_A^2+\sigma_B^2}+\frac{(A_y-B_y)^2}{\sigma_A^2+\sigma_B^2}\right) = \chi_2'^2\left(\frac{d^2_{\text{true}}}{\sigma_A^2+\sigma_B^2}\right). \tag{S4}$$

Fig. S1(a-b) show examples of the distributions of the normalized observed distance.

From Fig. S1, we see that as localization precision $\sigma_i$ increases, the mean of $d_{\text{obs}}/\sqrt{\sigma_A^2+\sigma_B^2}$ decreases. As $d_{\text{true}}$ increases, the mean of $d_{\text{obs}}/\sqrt{\sigma_A^2+\sigma_B^2}$ also increases, and the distribution starts to resemble a Gaussian. For any fixed $\sigma_A$ and $\sigma_B$, the monotonic dependence of $\mathbb{E}[d_{\text{obs}}/\sqrt{\sigma_A^2+\sigma_B^2}]$ on the normalized true distance (Fig. S1(c)) can be obtained using the exact formula for the mean of a non-central chi distribution (3). For completeness: the mean of $\chi_k'(\lambda)$ is given by

$$\mathbb{E}\chi_k'(\lambda) = \sqrt{2}\exp\left(-\frac{\lambda}{2}\right)\frac{\Gamma(\frac{k+1}{2})}{\Gamma(\frac{k}{2})}\text{hyp1f1}\left(\frac{k+1}{2},\frac{k}{2},\frac{\lambda}{2}\right)$$

where $\Gamma(z)$ is the gamma function,

$$\Gamma(z) = \int_0^\infty t^{z-1}e^{-t}\,dt,$$

and hyp1f1 is the confluent hypergeometric function 1F1,

$$\text{hyp1f1}(a,b,x) = \sum_{j=0}^{\infty}\frac{\Gamma(a+j)/\Gamma(a)}{\Gamma(b+j)/\Gamma(b)j!}x^j.$$

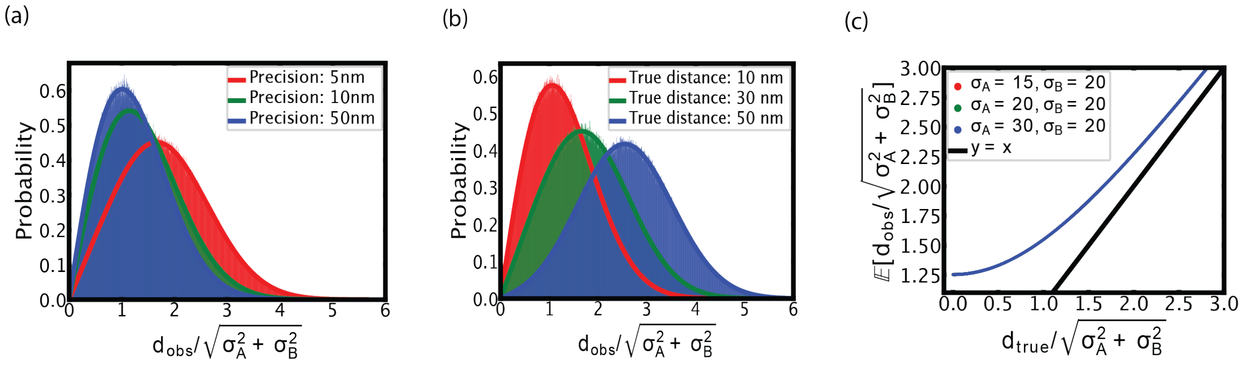

**Figure S1.** Theoretical distributions and histograms ($10^6$ sampled points) of $d_{\text{obs}}/\sqrt{\sigma_A^2+\sigma_B^2}$ simulated using Eq. (S4) for (a) $d_{\text{true}} = 10$ nm and localization precisions $\sigma_A = \sigma_B = 5, 10, 50$ nm, as well as (b) $\sigma_A = \sigma_B = 15$ nm and $d_{\text{true}} = 10, 30, 50$ nm. The simulated histograms align with the theoretical non-central chi-distributions characterized by a degree of freedom $k = 2$. (c) The mean of the random variable $d_{\text{obs}}/\sqrt{\sigma_A^2+\sigma_B^2}$ is independent of $\sigma_A$ and $\sigma_B$.

### Supplementary Note 2: Estimating the proximity probability of a pair

In this section, we describe a procedure to estimate the proximity probability $P_{\text{prox}}$ in Eq. (2) of the main text. As in the previous section, suppose emitters $A$ and $B$ are located at $(A_x, A_y)$ and $(B_x, B_y)$ respectively, with observed positions at $O_A$, $O_B$, and localization precisions $\sigma_A$, $\sigma_B$.

From Eq. (S2) above, let

$$Z_x = \frac{(O_{Ax} - O_{Bx}) - (A_x - B_x)}{\sqrt{\sigma_A^2 + \sigma_B^2}} \quad \text{and} \quad Z_y = \frac{(O_{Ay} - O_{By}) - (A_y - B_y)}{\sqrt{\sigma_A^2 + \sigma_B^2}},$$

so that $Z_x, Z_y \sim \mathcal{N}(0,1)$. Then the proximity probability $P_{\text{prox}}$ is given by

$$\begin{aligned}
&\mathbb{P}(d_{\text{lower}} \le d_{\text{true}} \le d_{\text{upper}} \,|\, d_{\text{obs}}, \sigma_A, \sigma_B) \\
&= \mathbb{P}(d_{\text{lower}}^2 \le |A_x - B_x|^2 + |A_y - B_y|^2 \le d_{\text{upper}}^2 \,|\, d_{\text{obs}}, \sigma_A, \sigma_B) \\
&= \mathbb{P}(d_{\text{lower}}^2 \le (O_{Ax} - O_{Bx} - \sqrt{\sigma_A^2+\sigma_B^2}\,Z_x)^2 + (O_{Ay} - O_{By} - \sqrt{\sigma_A^2+\sigma_B^2}\,Z_y)^2 \le d_{\text{upper}}^2). \qquad \textbf{(S5)}
\end{aligned}$$

The random variables in Eq. (S5) are $Z_x$ and $Z_y$. We can estimate $P_{\text{prox}}$ by generating many $Z_x, Z_y$, and counting how frequently the event in Eq. (S5) occurs. This process, whose pseudocode can be found in Algorithm 1, is computationally inexpensive, because it involves low-dimensional sampling. Typically $N = 10^5$ trials is sufficient to obtain a reliable estimate of the proximity probability.

**Algorithm 1** Monte Carlo procedure to estimate the proximity probability of two emitters

**Input**: $\sigma_A$, $\sigma_B$, $O_A$, $O_B$, $d_{\text{lower}}$, $d_{\text{upper}}$, number of MC trials $N$
**Output**: Proximity probability $P_{\text{prox}}$
$m \leftarrow 0$
  **for** $i \leftarrow 1$ **to** $N$ **do**
    Sample $Z_x \sim \mathcal{N}(0,1)$, $Z_y \sim \mathcal{N}(0,1)$
    **if** $d_{\text{lower}}^2 \le (O_{Ax} - O_{Bx} - \sqrt{\sigma_A^2+\sigma_B^2}\,Z_x)^2 + (O_{Ay} - O_{By} - \sqrt{\sigma_A^2+\sigma_B^2}\,Z_y)^2 \le d_{\text{upper}}^2$ **then**
      $m \leftarrow m + 1$
    **end if**
  **end for**
**return** $P_{\text{prox}} \approx \frac{m}{N}$

Fig. S2 shows the effect of observation parameters on estimating $P_{\text{prox}}$. In (a-c), the left panels show the true positions of the two emitters, with the black bar indicating $d_{\text{obs}}$; the right panels show the Gaussians $\mathcal{N}(O_A - O_B, \sigma_A^2 + \sigma_B^2)$, and green rings marking the annulus $0 = d_{\text{lower}} \le r \le d_{\text{upper}}$. The proximity probability is maximized when most of the mass of $\mathcal{N}(O_A - O_B, \sigma_A^2 + \sigma_B^2)$ lies within the annulus (Fig. S2(a,d)). The examples showcased in Fig. S2 are computed as follows, with all units in nm:

$$\mathbb{P}(0 \leq d_{\text{true}} \leq 20 | d_{\text{obs}} = 5, \sigma_A = \sigma_B = 5) \approx 96\%,$$
$$\mathbb{P}(0 \leq d_{\text{true}} \leq 25 | d_{\text{obs}} = 10, \sigma_A = 15, \sigma_B = 10) \approx 56\%,$$
$$\mathbb{P}(0 \leq d_{\text{true}} \leq 25 | d_{\text{obs}} = 50, \sigma_A = 15, \sigma_B = 25) \approx 13\%.$$

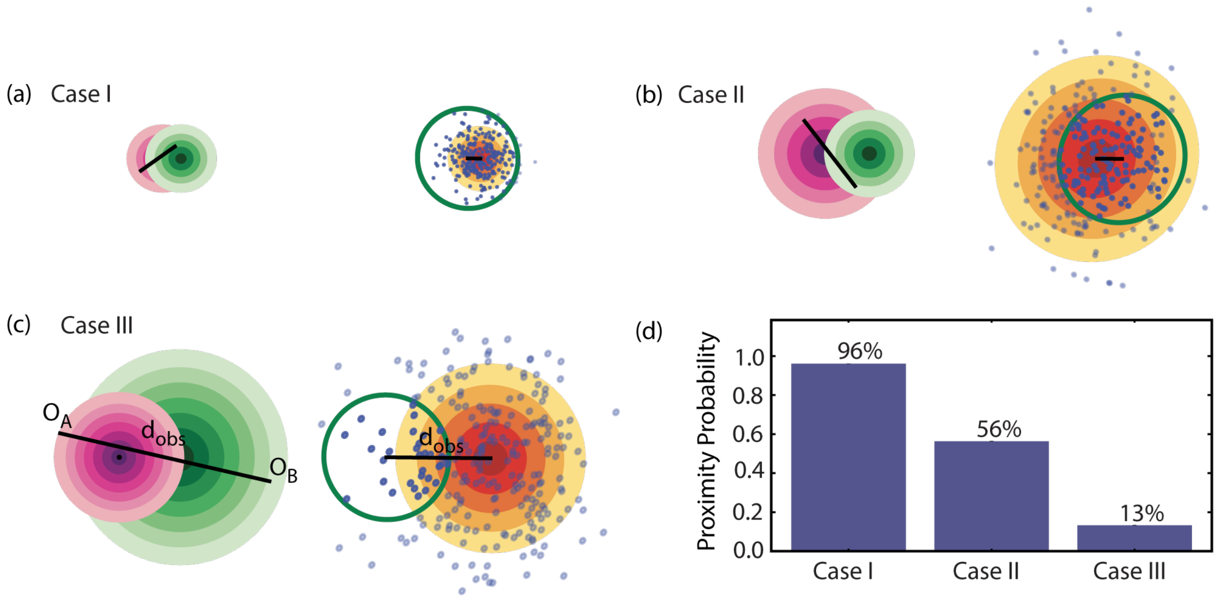

**Figure S2.** Monte Carlo procedure for estimating the proximity probability $P_{\text{prox}}$ in three scenarios (not drawn to scale). (a-c) Pink (respectively green) rings represent the distribution for $O_A \sim \mathcal{N}((A_x, A_y), \sigma_A^2)$ (respectively $O_B \sim \mathcal{N}((B_x, B_y), \sigma_B^2)$); black bars mark observed distances $d_{\text{obs}}$. Yellow rings represent $\mathcal{N}(O_A - O_B, \sigma_A^2 + \sigma_B^2)$, from which points were drawn. The fraction of points that land inside the green ring estimates $P_{\text{prox}}$. (d) The approximated $P_{\text{prox}}$ is largest when the localization uncertainties are small and the observed displacement lies in the annulus.

### Supplementary Note 3: Data-driven selection of a threshold parameter for $d_{\text{obs}}$

Intuitively, if $A$ and $B$ are extremely far away, i.e., $d_{\text{obs}} \gg d_{\text{true}}$, we would not expect them to pair up. By considering only emitters that are reasonably close by, we cut down the computational time for Graph Matching Optimization (GMO). In this section, we quantify this threshold, and provide a heuristic for how to select the threshold parameter for $d_{\text{obs}}$ based on the statistics of the localizations in a SMLM image.

In principle, GMO selects a matching from all possible pairs. However, using all possible pairs to construct the bipartite graph would be computationally expensive, and with diminishing return for the performance of GMO. For example, with $N_A$ molecules $A$ and $N_B$ molecules $B$, there can be up to $N_A N_B / 2$ possible pairs. Selecting the maximum weight matching from the input bipartite graph, as implemented in NetworkX (4; 5), is computationally expensive, with time complexity that scales as $O(n^3)$, where $n$ denotes the number of nodes. Thus, when building the bipartite graph for GMO, we only consider pairs that are within a threshold distance away from each other. We do this by cutting out the thin tail of the distribution in Eq. (S4) and considering only pairs where $d_{\text{obs}}/\sqrt{\sigma_A^2 + \sigma_B^2} \leq C$ for some thresholding parameter $C$.

This truncation of the tail has minimal effect on the distribution. For example, for the distributions in Fig. S1(a), choosing $C = 4$ captures the bulk of the distributions:

$$\mathbb{P}(d_{\text{obs}}/\sqrt{\sigma_A^2 + \sigma_B^2} \leq 4 | d_{\text{true}} = 10, \sigma_A = \sigma_B = 5) = 99.13\%,$$
$$\mathbb{P}(d_{\text{obs}}/\sqrt{\sigma_A^2 + \sigma_B^2} \leq 4 | d_{\text{true}} = 10, \sigma_A = \sigma_B = 10) = 99.87\%,$$
$$\mathbb{P}(d_{\text{obs}}/\sqrt{\sigma_A^2 + \sigma_B^2} \leq 4 | d_{\text{true}} = 10, \sigma_A = \sigma_B = 50) = 99.96\%.$$

Similarly, for the distributions in Fig. S1(b):

$$\mathbb{P}(d_{\text{obs}}/\sqrt{\sigma_A^2+\sigma_B^2} \le 4 \,|\, d_{\text{true}} = 10, \sigma_A = \sigma_B = 15) = 99.93\%,$$
$$\mathbb{P}(d_{\text{obs}}/\sqrt{\sigma_A^2+\sigma_B^2} \le 4 \,|\, d_{\text{true}} = 30, \sigma_A = \sigma_B = 15) = 99.13\%,$$
$$\mathbb{P}(d_{\text{obs}}/\sqrt{\sigma_A^2+\sigma_B^2} \le 4 \,|\, d_{\text{true}} = 50, \sigma_A = \sigma_B = 15) = 93.05\%.$$

Clearly, we cannot blindly choose $C = 4$ for any SMLM image. To choose $C$, one must first estimate $d_{\text{true}}$ for the two interacting molecule partners. Such an estimate may be obtained from fitting the distributions in Eq. (S1) from experiments such as iterative localization of the identical molecular complex (6; 2; 7). Next, one would plot the distribution of localization precision values of the SMLM image, and select a threshold $\sigma_*$ such that it captures most of the precision values, e.g., $\sigma_i \le \sigma_*$ captures the 90th percentile of all $\sigma_A, \sigma_B$ from the dataset. One can then simulate the distribution of $d_{\text{obs}}/\sqrt{2\sigma_*^2}$ according to Eq. (S4). From the resulting distribution similar to those in Fig. S1(a-b), $C$ can be chosen such that $\{d_{\text{obs}}/\sqrt{\sigma_A^2+\sigma_B^2} \le C\}$ captures the bulk of the distribution.

In principle, a larger value of $C$ would capture more possible pairings. However, small increases in $C$ might significantly increase the computational time due to the increased density of the graph. Choices must be made based on the dataset being analyzed, and the goal of the analysis.

#### Supplementary Note 4: Estimating the true number of molecular couplings and the number of pairings from transient proximity

Because SMLM captures the observed positions of emitters at a fixed time, we cannot distinguish between molecular coupling and transient proximity. For example, if the SMLM image is a snapshot of Fig. S3 at $t_2$, we cannot distinguish between the two scenarios. We devised a process we call Iterative Monte Carlo Estimation of Molecular Couplings and Background Pairings (iMEC) as part of our colocalization algorithm, to estimate how many pairs are true couplings in a given SMLM dataset, and how many are due to transient proximity. More precisely, given $N_A$ and $N_B$ emitters, of which there are $N^*_{\text{pairs}}$ pairings identified by GMO, Algorithm 2 estimates how many of the $N^*_{\text{pairs}}$ pairings are transient, in the sense of Fig. S3(b). Thus, the number of true molecular couplings is what remains after subtracting the transient pairs.

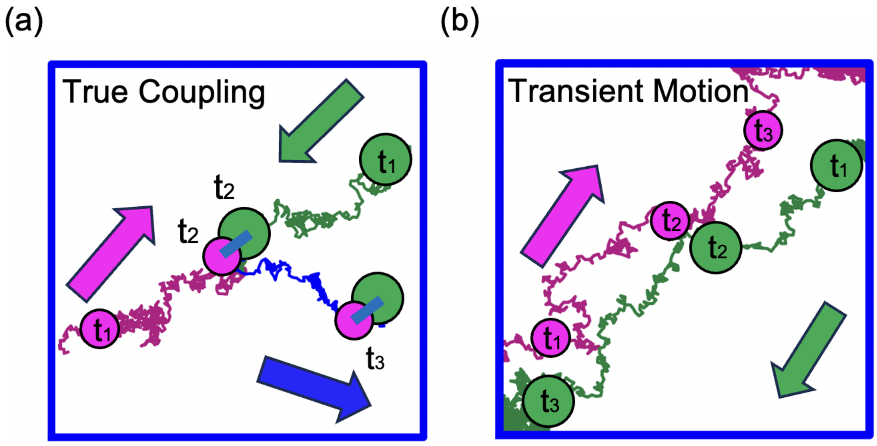

**Figure S3.** Stochastic Brownian trajectories of two interacting partners. (a) Coupling occurs between two molecules at time $t_2$, after which they proceed together until $t_3$ along the trajectory in blue. (b) Two molecules experience transient proximity during diffusion. A measurement only at time $t_2$ cannot distinguish between the two scenarios.

We assume a system where $A$ and $B$ diffuse freely. If a molecule of $A$ and a molecule of $B$ are sufficiently close, they are capable of binding (Fig. S3(a)). However, if the system does not pass the activation energy barrier for the binding reaction to occur, the two molecules may simply diffuse away at a later time (Fig. S3(b)). A comprehensive stochastic model elucidating the pivotal role of colocalization in determining the probability of protein-protein interactions is detailed in the works (8; 9) by Batada et al..

In this work, we take a slightly different approach. In iMEC (Algorithm 2), we successively estimate the number of pairings among molecules that we assume cannot interact, with a smaller number of molecules at the start of each iteration. The inputs to Algorithm 2 are the number of signals corresponding to $A$ ($N_A$), the number of signals corresponding to $B$ ($N_B$), the number of pairs identified by GMO ($N^*_{\text{pairs}}$), and the area of the region being analyzed ($S$) . In addition we specify the number of iterations per trial ($T$), and the number of trials ($M$).

In any given trial, for the first iteration, we assume all $N_A$ and $N_B$ molecules are non-interacting ($N^0_{\text{coupled}} = 0$), and estimate the number of pairs $N^1_{\text{bg}}$. All pairs identified at this step are considered background pairs since we assumed non-interacting particles. Then we set $N^1_{\text{coupled}} = N^*_{\text{pairs}} - N^1_{\text{bg}}$. For the second iteration, since we assume there are at least $N^1_{\text{coupled}}$ coupled pairs, we repeat the process by assuming that the remaining $N_A - N^1_{\text{coupled}}$ molecules of $A$ and $N_B - N^1_{\text{coupled}}$ molecules of $B$ are non-interacting. After each iteration, we update the estimated number of background pairs $N^i_{\text{bg}}$, and thus obtain an estimated number of true couplings $N^i_{\text{coupled}} = N^*_{\text{pairs}} - N^i_{\text{bg}}$. We stop the trial after $T$ iterations, and obtain estimates of $N^T_{\text{bg}}$ and $N^T_{\text{coupled}}$. The output of Algorithm 2 is the average over $M$ trials: $N^*_{\text{coupled}} \approx \langle N^T_{\text{coupled}} \rangle$.

An example of $\langle N^t_{\text{bg}} \rangle$ and $\langle N^t_{\text{coupled}} \rangle$ is shown in Fig. **??** of the main text. In the main text, we suppress the notation for multiple MC trials for simplicity, i.e., $N^T_{\text{bg}}$ in the main text is in actuality $\langle N^T_{\text{bg}} \rangle$, and likewise for the number of molecular couplings.

In principle, it is possible that $N^i_{\text{bg}} > N^*_{\text{pairs}}$ at the end of an iteration, so $N^i_{\text{coupled}} < 0$ by definition. This reflects the situation where there are more background pairs expected than those identified by GMO. In such a case, the next iteration starts with $N_A - N^i_{\text{coupled}}$ and $N_B - N^i_{\text{coupled}}$ molecules of $A$ and $B$ respectively. Note that here we instantiate more molecules than in the dataset. If this occurs at the last iteration, we set $N^T_{\text{coupled}} = 0$. However, this is unlikely to occur in a dataset for which molecular couplings are expected.

**Algorithm 2** Iterative Monte Carlo procedure to estimate the number of molecular couplings.

**Input**: $N_A$, $N_B$, $N^*_{\text{pairs}}$, area $S$, number of iterations $T$ per trial, number of MC trials $M$.
**Output**: $\langle N^T_{\text{bg}} \rangle$, $\langle N^T_{\text{coupled}} \rangle$
$S_{\text{bg}} \leftarrow 0$
$S_{\text{coupled}} \leftarrow 0$
  **for** $m \leftarrow 1$ **to** $M$ **do**
    $N^0_{\text{coupled}} \leftarrow 0$
    **for** $i \leftarrow 1$ **to** $T$ **do**
      Simulate $N_A - N^{i-1}_{\text{coupled}}$ and $N_B - N^{i-1}_{\text{coupled}}$ particles over area $S$ uniformly random.
      Use pairing algorithm to find the most probable graph matching to estimate $N^i_{\text{bg}}$.
      $N^i_{\text{coupled}} \leftarrow N^*_{\text{pairs}} - N^i_{\text{bg}}$
    **end for**
    $S_{\text{bg}} \leftarrow S_{\text{bg}} + N^T_{\text{bg}}$
    $S_{\text{coupled}} \leftarrow S_{\text{coupled}} + N^T_{\text{coupled}}$
  **end for**
  $\langle N^T_{\text{bg}} \rangle \leftarrow S_{\text{bg}}/M$
  $\langle N^T_{\text{coupled}} \rangle \leftarrow S_{\text{coupled}}/M$
  **if** $\langle N^T_{\text{coupled}} \rangle < 0$ **then**
    Set $\langle N^T_{\text{coupled}} \rangle = 0$
  **end if**
**return** $\langle N^T_{\text{bg}} \rangle$, $\langle N^T_{\text{coupled}} \rangle$

### Supplementary Note 5: Pipeline to estimate the number of molecule couplings in an SMLM image

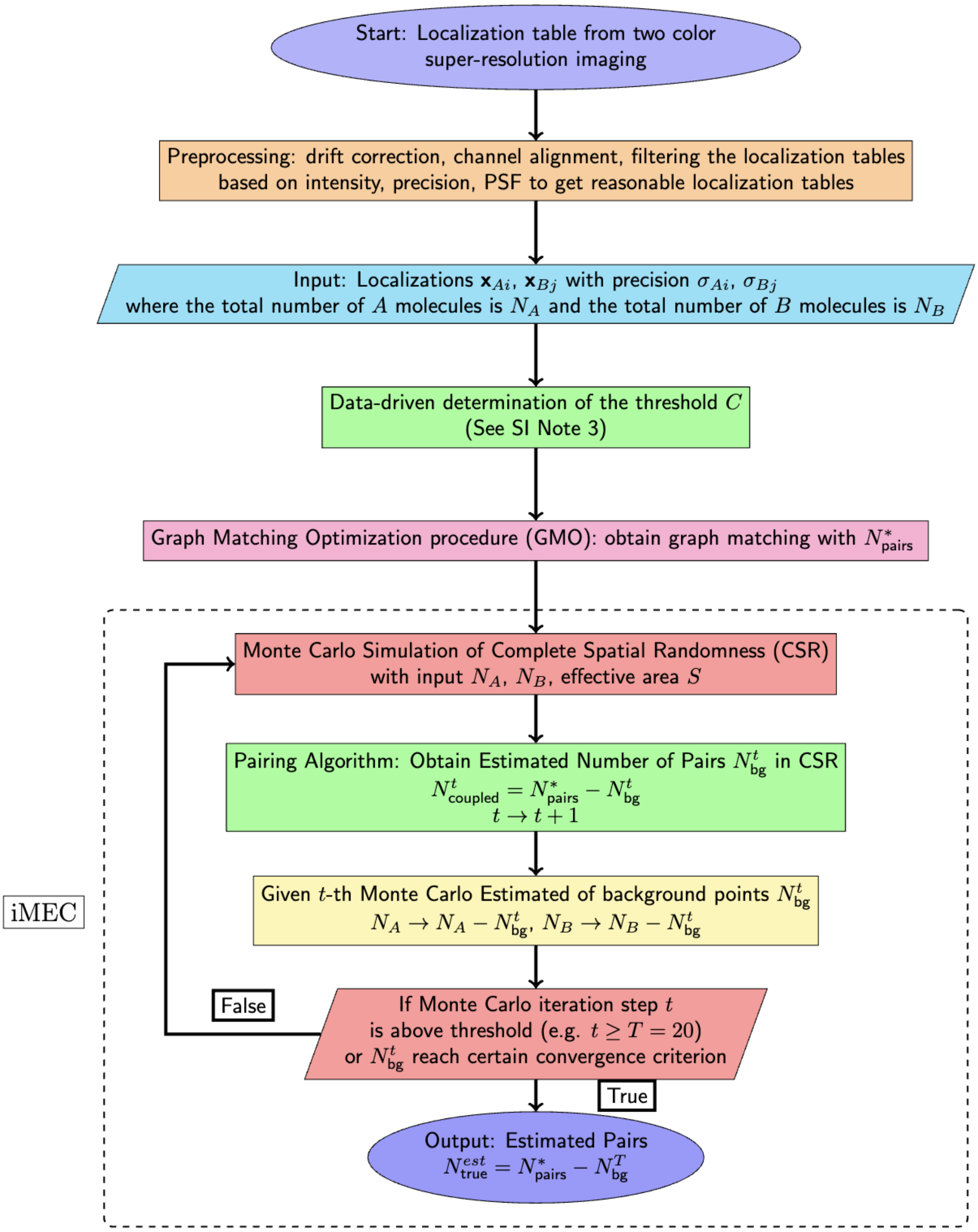

**Figure S4.** Full pipeline to detect molecular pairings and count the number of true couplings in a given SMLM localization table. The core components include a graph matching optimization procedure (GMO) and an iterative Monte Carlo estimation of molecular couplings (iMEC).

### Supplementary Note 6: Generating simulated SMLM datasets

Many of our results were based on simulated SMLM datasets. To generate a simulated dataset, we instantiated the positions of molecules $A$, $B$, and $AB$ either from a uniform distribution, or from a stochastic reaction-diffusion simulation.

#### A. Generating two-channel signals

To generate the localizations in the two channels from the positions of $A$, $B$, and $AB$, we kept the localizations for $A$ and $B$, and split each $AB$ into two localizations. For each $AB$ located at $(x, y)$, we sampled from the uniform distribution a stochastic distance $d_{\text{obs}} \sim \text{Unif}(0, 10\,\text{nm})$, and an angle $\theta \sim \text{Unif}(0, 2\pi)$. We placed signals corresponding to $A$ and $B$ at

$$\left(x + \frac{d_{\text{obs}}}{2}\cos\theta,\, y + \frac{d_{\text{obs}}}{2}\sin\theta\right) \quad \text{and} \quad \left(x - \frac{d_{\text{obs}}}{2}\cos\theta,\, y - \frac{d_{\text{obs}}}{2}\sin\theta\right)$$

respectively. For the localization uncertainty of each signal, we either set it as a constant (e.g., for Fig. 4 of the main text; see Tables S3), or drew from the distribution in Fig. S6.

#### B. Stochastic simulation

The stochastic simulations reported in Fig. 6 of the main text were generated using a particle-based reaction–diffusion model, implemented using the package ReaDDY, version 2.0.12 (10).

We set the simulation box to be $\sqrt{\text{Area}} \times \sqrt{\text{Area}} \times 1$ nm, with periodic boundaries. This effectively simulates the system as if it is 2-dimensional for Area $\gg 1$. Diffusion is modeled by $\frac{d\vec{x}_i(t)}{dt} = \sqrt{2D_i}\vec{\xi}_i(t)$, where $\vec{x}_i(t)$ is the position of particle $i$, with diffusion coefficient $D_i$, and $\{\vec{\xi}_i\}_i$ are independent Gaussian random variables.

Each specified reaction occurs with probability $p(\lambda; dt) = 1 - e^{-\lambda dt}$, where $\lambda$ plays a similar role to a rate constant. For a reaction involving a single reactant, e.g., $AB \to A + B$, the propensity constant is given by the macroscopic rate constant: $\lambda_{\text{off}} = k_{\text{off}}$. For binary reactions, e.g., $A + B \to AB$, a reaction would occur with probability $p(\lambda; dt)$ only if the reacting molecules, $A$ and $B$ in this case, are no more than $d_{\text{educt}}$ apart, where the educt distance $d_{\text{educt}}$ is a chosen parameter. For this simulation, we chose $d_{\text{educt}} = 30$ nm. Note that for binary reactions, $k_{\text{on}} \neq \lambda_{\text{on}}$ in general. For $d_{\text{educt}} = 30$ nm, the relationship for $\lambda_{\text{on}} \leq 25\,\text{s}^{-1}$ was numerically determined to be

$$k_{\text{on}} = 2.72 \times 10^{-3}\lambda_{\text{on}} + 4.54 \times 10^{-4},$$

as shown in Fig. S5. For $\lambda_{\text{on}} = 20\,\text{s}^{-1}$, the corresponding rate constant is $k_{\text{on}} = 0.0549$ µm$^2$molecule$^{-1}$s$^{-1}$.

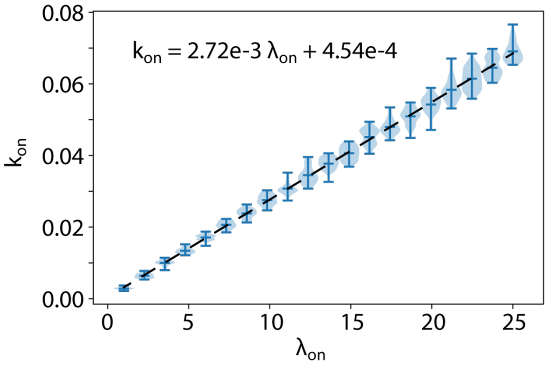

**Figure S5.** Conversion between the rate constant $k_{\text{on}}$ and the propensity constant $\lambda_{\text{on}}$ for $A + B \to AB$, at $d_{\text{educt}} = 30$ nm.

We simulated the system with initially only $A$ and $B$, but no $AB$. At the densities stated in Fig. 6(c) (pink 30 $A$/µm$^2$, 20 $B$/µm$^2$; blue 20 $A$/µm$^2$, 20 $B$/µm$^2$; yellow 10 $A$/µm$^2$, 20 $B$/µm$^2$), the particles were uniformly distributed in the simulation box. We ran the simulation, and extracted the positions of $A$, $B$, and $AB$ at various time points. The parameters for the stochastic simulations are presented in Table S1.

**Table S1.** Stochastic Simulation Parameters in Fig. 6

| | |
|---|---|
| Timestep $dt$ | $0.5 \times 10^{-3}$ s |
| Total Simulation Time | 3 s |
| Area | 100 $\mu$m$^2$ |
| $\lambda_{\text{on}}$ | 20 s$^{-1}$ |
| $\lambda_{\text{off}}$ | 1 s$^{-1}$ |
| $d_{\text{educt}}$ | 30 nm |
| Diffusion coefficient $D_A$ | 0.3 $\mu$m$^2$/s |
| Diffusion coefficient $D_B$ | 0.3 $\mu$m$^2$/s |
| Diffusion coefficient $D_{AB}$ | 0.01 $\mu$m$^2$/s |
| Densities of $(A, B)$ | (30,20), (20,20) or (10,20) molecules/$\mu$m$^2$ |

##### C. An experimental distribution of localization precisions

The average experimental precision in our datasets typically ranged from 15 to 30 nm. In several of our simulated SMLM datasets, we used localization precision values drawn from the experimental distribution in Fig. S6, with $\sigma_i \leq 40\,$nm to model localizations with moderate to high signal-to-noise ratios. We found that the localization precision distributions between different cells (under the same experimental conditions) were remarkably similar. Fig. S6 is representative of the typical distributions observed; experimental protocol can be found in Supplementary Note 9.

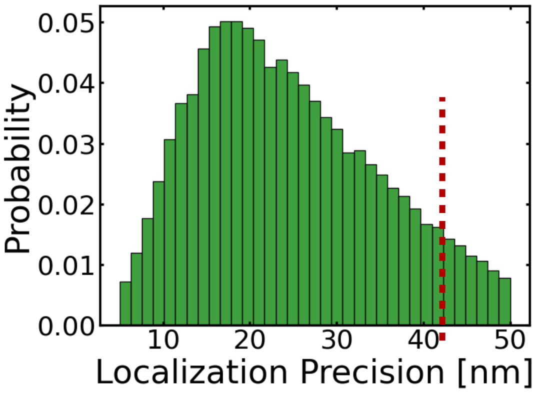

**Figure S6.** An experimental localization precision distribution with the peak at approximately 20 nm. The cutoff localization precision used in all simulations was 40 nm.

#### Supplementary Note 7: Parameters used for various analyses

For Figures 3-6 in the main text, to produce the results, we ran GMO and iMEC. This section consists of tables of parameters used in the analysis.

**Table S2.** Simulation Parameters for Fig. 3(b-c)

| | |
|---|---|
| Area $S$ | 100 $\mu$m$^2$ |
| Number of AB | 1545 or 1100 |
| Number of background A $N_A$ | 455 or 900 |
| Number of background B $N_B$ | 455 or 900 |
| True distance $d_{\text{true}}$ | Sampled from Unif(0,10 nm) |
| Localization Precision $\sigma_A, \sigma_B$ | Sampled from Fig. S6 |
| $d_{\text{lower}}$ | 0 nm |
| $d_{\text{upper}}$ | 20 nm |
| Search distance factor threshold $C$ from SN 3 | 4 |
| Number of sampling points to estimate probability $N$ from Algorithm 1 | $10^5$ |
| Number of iterations $T$ to estimate $N^*_{\text{bg}}$ from Algorithm 2 | 15 |
| Number of MC trials $M$ in each estimation of $N^i_{\text{bg}}$ in Algorithm 2 | 10 |

**Table S3.** Simulation Parameters in Fig. 4

| | Fig. 4(a) | Fig. 4(b) |
|---|---|---|
| Density $\rho$ | $\rho =$ 5, 10 or 20 (see plot) | Varying (see $x$-axis) |
| Area $S$ | $S =$ 100 $\mu$m$^2$ | 100 $\mu$m$^2$ |
| Number of AB | $\rho S/2$ | $\rho S/2$ |
| Number of background A | $\rho S/2$ | $\rho S/2$ |
| Number of background B | $\rho S/2$ | $\rho S/2$ |
| True distance $d_{\text{true}}$ | Sampled from Unif(0,10 nm) | Sampled from Unif(0,10 nm) |
| Localization Precision $\sigma_A$, $\sigma_B$ | Varying (see $x$-axis) | $\sigma_A = \sigma_B =$ 10, 20 or 30 nm (see plot) |
| $d_{\text{lower}}$ | 0 nm | 0 nm |
| $d_{\text{upper}}$ | 20 nm | 20 nm |
| Search distance factor threshold $C$ from SN 3 | 4 | 4 |
| Number of sampling points $N$ to estimate probability from Algorithm 1 | $10^5$ | $10^5$ |
| Number of iterations $T$ to estimate $N^*_{\text{bg}}$ from Algorithm 2 | 25 | 25 |
| Number of MC trials $M$ in each estimation of $N^i_{\text{bg}}$ in Algorithm 2 | 10 | 10 |

**Table S4.** Simulation Parameters in Fig. 5(d)

| | |
|---|---|
| Percentage of couplings $c$ | $c =$ 0%, 6% or 21% |
| Density $\rho_A$ | $\rho_A =$ 5 $\mu$m$^{-2}$ |
| Density $\rho_B$ | $\rho_B =$ 5, 10, 15 $\mu$m$^{-2}$ |
| Area $S$ | $S =$ 100 $\mu$m$^2$ |
| Number of AB | $\rho_A S c$ |
| Number of background A | $\rho_A S(1-c)$ |
| Number of background B | $\rho_B S(1-c)$ |
| $K_{\text{eq}}$ | $\min(\rho_A, \rho_B)c/(1-c)^2 \rho_A \rho_B$ |
| True distance $d_{\text{true}}$ | Sampled from Unif(0,10 nm) |
| Localization Precision $\sigma_A$, $\sigma_B$ | Sampled from Fig. S6 |
| $d_{\text{lower}}$ | 0 nm |
| $d_{\text{upper}}$ | 20 nm |
| Search distance factor threshold $C$ from SN 3 | 4 |
| Number of sampling points to estimate probability $N$ from Algorithm 1 | $10^5$ |
| Number of iterations $T$ to estimate $N^*_{\text{bg}}$ from Algorithm 2 | 25 |
| Number of MC trials $M$ in each estimation of $N^i_{\text{bg}}$ in Algorithm 2 | 10 |

**Table S5.** Simulation Parameters in Fig. 6(c-d)

| | |
|---|---|
| Density $\rho$ | $\rho_A = 30, 20, 10$, $\rho_B = 20$ µm$^{-2}$ |
| Area $S$ | 100 $\mu$m$^2$ |
| Number of AB at time $t$ | From stochastic simulation |
| Number of background A at time $t$ | From stochastic simulation |
| Number of background B at time $t$ | From stochastic simulation |
| True distance $d_{\text{true}}$ | Sampled from Unif(0,10 nm) |
| Localization Precision $\sigma_A$, $\sigma_B$ | Sampled from Fig. S6 |
| $d_{\text{lower}}$ | 0 nm |
| $d_{\text{upper}}$ | 20 nm |
| Search distance factor threshold $C$ from SN 3 | 4 |
| Number of sampling points to estimate probability $N$ from Algorithm 1 | $10^5$ |
| Number of iterations $T$ to estimate $N^*_{\text{bg}}$ from Algorithm 2 | 25 |
| Number of MC trials $M$ in each estimation of $N^i_{\text{bg}}$ in Algorithm 2 | 10 |

### Supplementary Note 8: Comparing GMO's performance against an alternative method

Most colocalization methods focus on evaluating spatial correlations between the two channels, and do not directly address colocalization between two specific signals (11; 12; 13; 14; 15). In order to compare our method (GMO) against an alternative, we designed an algorithm based on the ideas in (16). We called this the *Minimal Pairwise Distance Method* (MinDist), as it aims to minimize distances between pairs.

#### A. Colocalization by the Minimal Pairwise Distance Method

MinDist sorts all pairwise distances $d_{\text{obs}}$ (within a threshold $d_{\text{thres}}$), and selects colocalized pairs starting with the smallest distances. After each pair has been selected, their corresponding signals are no longer available for further colocalization, so we delete their pairwise distances to any other localizations from the list. The pseudo-code is provided in Algorithm 3.

**Algorithm 3** Minimal Pairwise Distance Method (MinDist)

**Input**: $d_{\text{thres}}$, list of observed positions $\{O_{A_1},\ldots,O_{A_N}\}$, $\{O_{B_1},\ldots,O_{B_M}\}$
**Output**: list of pairs $C = \{(i,j)\}$ of putative colocalizations

```
I ← [ ]
D ← [ ]
C ← [ ]
for n ← 1 to N, m ← 1 to M do
    if d_nm = dist(O_An, O_Bm) < d_thres then
        Add d_nm to D
        Add (n,m) to I
    end if
    Sort I and D by ascending order of D
    while I is not empty do
        Add the first entry (n,m) ∈ I to C
        Delete all entries (n,j) from I and the corresponding entries d_nj from D
        Delete all entries (i,m) from I and the corresponding entries d_im from D
    end while
end for
return C
```

Although MinDist is very intuitive, it has two obvious drawbacks. 1) It does not take into account localization precision values, and 2) it is unclear how $d_{\text{thres}}$ should be chosen. The purpose of $d_{\text{thres}}$ is to speed up computation. As $d_{\text{thres}} \to \infty$, all possible pairs are being considered. For finite $d_{\text{thres}}$, as it increases, more putative pairs are being considered, leading to increases in both true and false positives.

#### B. Parameters used in GMO and MinDist

We compared GMO and MinDist in two scenarios: short interaction distance ($15 \leq d_{\text{true}} \leq 25$ nm) and long interaction distance ($55 \leq d_{\text{true}} \leq 65$ nm). Parameters used in each scenarios are presented in Table S6.

As noted earlier, a larger value of $d_{\text{thres}}$ in MinDist in general increases both the number of true positives (TP) and false positives (FP). To ensure a fair comparison, we set $d_{\text{thres}}$ so that FP for GMO is nearly identical to that of MinDist (see right-panels of Figs. S7(a) and S8(a)).

**Table S6.** Parameters used in the comparison of GMO to MinDist

| | **Small interaction distance** | **Large interaction distance** |
|---|---|---|
| Density $\rho$ | Varying (see $x$-axis of Fig. S7) | Varying (see $x$-axis of Fig. S7) |
| Area $S$ | $S = 100$ | 100 |
| Number of AB | $\rho S$ | $\rho S$ |
| Number of background A | 0 | 0 |
| Number of background B | 0 | 0 |
| True distance $d_{\text{true}}$ | Sampled from Unif(15, 25 nm) | Sampled from Unif(55, 65 nm) |
| Localization Precision $\sigma_A, \sigma_B$ | Sampled from Fig. S6 | Sampled from Fig. S6 |
| $d_{\text{lower}}$ | 10 nm | 45 nm |
| $d_{\text{upper}}$ | 30 nm | 75 nm |
| Search distance factor threshold $C$ from SN 3 | 4 | 4 |
| Number of sampling points to estimate probability | $10^5$ | $10^5$ |
| $d_{\text{thres}}$ in minimal pairwise distance method | 85 nm | 90 nm |

#### C. Results from comparing GMO to MinDist

We evaluated the performance of the two methods by comparing the number of true positives per micron (TP), the number of false positives per micron (FP), the TP ratio of GMO to MinDist, recall, and precision, where

$$\text{Recall} = \frac{\text{TP}}{\text{TP} + \text{FN}} \quad \text{and} \quad \text{Precision} = \frac{\text{TP}}{\text{TP} + \text{FP}}.$$

In all these metrics, GMO outperformed MinDist.

Consider first the case of short interaction distance, where we sampled $d_{\text{true}}$ from $[15, 25]$ nm uniformly. Fig. S7(a) plots TP and FP for GMO and MinDist. We chose $d_{\text{thres}}$ to minimize the difference in FP. GMO clearly outperformed MinDist by at least 10%, as demonstrated by the ratio of TPs in Fig. S7(b). Moreover, this performance improved slightly as the density of localization signals increased. Fig. S7(c-d) show recall and precision of the two algorithms. GMO's recall was constantly $\sim$10% better than MinDist's, while precision was marginally better.

We also considered the scenario where the interaction distance was large. For example, (13) reported an interaction distance of $\sim$60 nm between PSD95 and Homer, which are involved in the functioning of synapses. For long interaction distance, we uniformly sampled $d_{\text{true}}$ from [55, 65] nm. Fig. S8(a) shows TP and FP for both GMO and MinDist, where again, $d_{\text{thres}}$ for MinDist was chosen to minimize the difference in FP's. Here GMO performed significantly better than MinDist at all densities, by $\sim$40-50% (Fig. S8(b)). In terms of recall, Fig. S8(c) shows that GMO outperformed MinDist by $\sim$25% across all densities, and GMO's precision was increasingly better than MinDist's at higher density (Fig. S8(d)).

At all interaction distances, GMO constantly outperformed MinDist both in the number of true positives discovered and recall, across all densities of localizations. The outperformance was more significant for longer interaction distances. This is unsurprising as MinDist prioritizes *shorter* pairwise distances for its colocalizations.

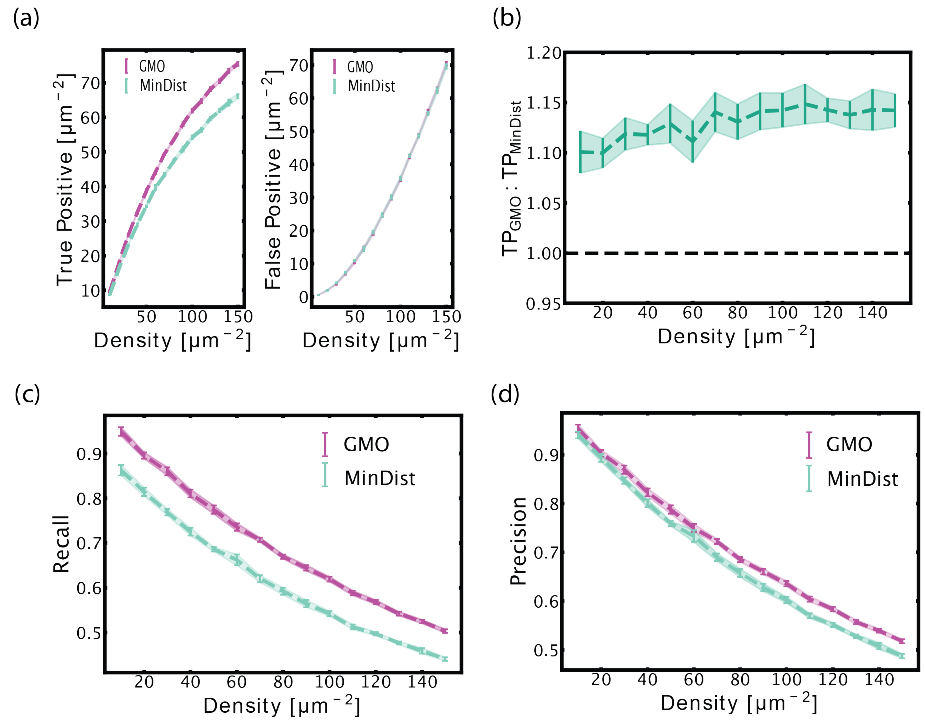

**Figure S7.** Performance of GMO versus MinDist at short interaction distance. (a) True Positives (TP) and False Positives (FP) discovered at different densities. (b) The ratio of TPs from the two methods, where $TP_{GMO}$:$TP_{MinDist} > 1$ means GMO outperforms MinDist at identifying TPs. In terms of (c) recall and (d) precision, GMO outperformed MinDist. Error bars indicate $\pm 1$ standard deviation over 10 simulated datasets.

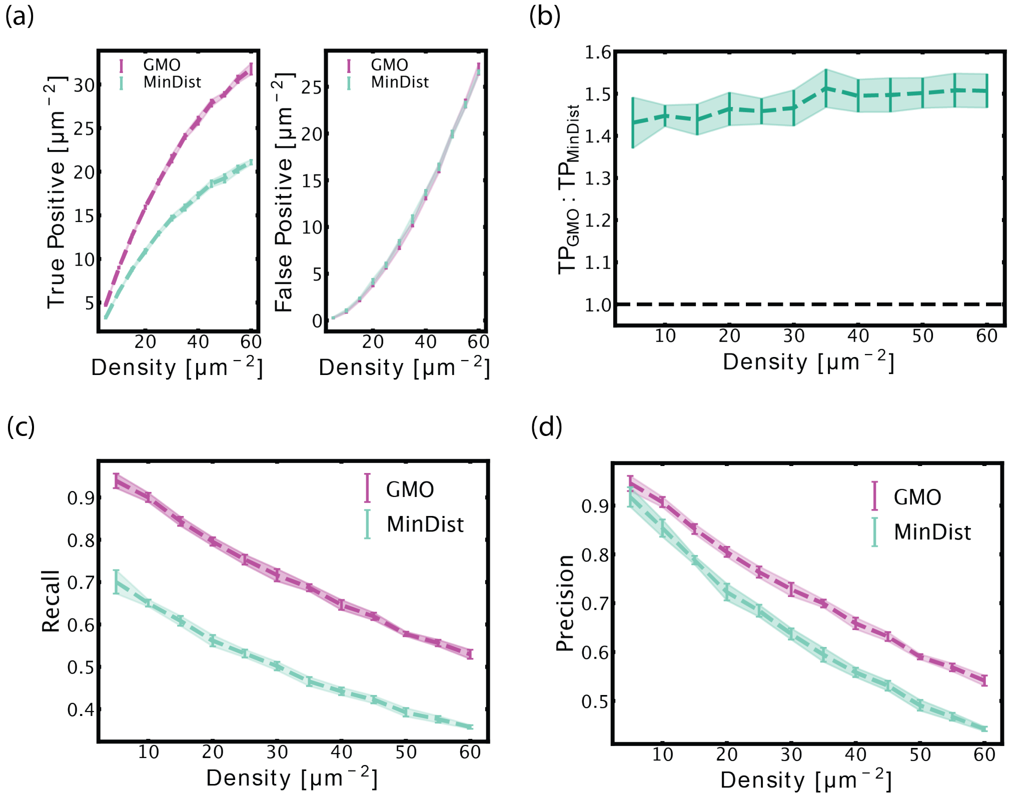

**Figure S8.** Performance of GMO versus MinDist at long interaction distance. (a) True Positives (TP) and False Positives discovered at different densities. (b) The ratio of TPs from the two method, where $TP_{GMO}$:$TP_{MinDist} > 1$ means GMO does better picking up TPs. In terms of (c) recall and (d) precision, GMO outperformed MinDist. Error bars indicate $\pm 1$ standard deviation over 10 simulated datasets.

### Supplementary Note 9: Experimental materials and methods

#### A. Cells and DNA constructs

***A.1. Cell lines.*** Two cell lines were used: Human Embryonic Kidney 293FT (HEK293FT) cells and HEK293 cells with a CRISPR/Cas9 knockout of the G-protein Gs (HEK293 ΔGs). HEK293 ΔGs cells were received as a gift from Dr. Asuka Inoue at Tohoku University (17; 18; 19).

***A.2. Cell culture and transfection.*** Cells were cultured in Dulbecco's Modification of Eagle's Medium (DMEM) with 4.5 g/L glucose, L-glutamine & sodium pyruvate (Corning CAT #10-013-CV). DMEM (500 mL) was supplemented with 50 mL of Fetal Bovine Serum (FBS) (VWR CAT #89510-186) and 5 mL of 100x Penicillin-Streptomycin solution (Corning CAT #30-002-Cl). Media was sterilized by filtration with a 0.2 µm PES bottle filter (CAT #10040-436). Culture media for live-cell confocal imaging was prepared the same way except that Phenol-Red-free FluoroBrite DMEM (Fisher Scientific CAT #A1896701) was used.

Cells were cultured in 25 $cm^2$, 75 $cm^2$, and 175 $cm^2$ treated culture flasks (VWR #10062-872, GenClone #25209, Corning #353136, VWR #10062-864) using standard sterile techniques. Passaging was done using a solution of 0.05% Trypsin + 0.53 mM EDTA in Phosphate-Buffered Saline (PBS) without calcium and magnesium (Corning CAT #21-040-CV). A 10x stock of this buffer was made by diluting 2.5% trypsin (ThermoFisher #15090046) and powdered EDTA (CalbioChem CAT #4050) to 0.5% trypsin + 5.3 mM EDTA, pH = 7.4. Stock solution was frozen in 1 mL aliquots at -20°C in 15 mL conical centrifuge tubes. Before use, the 1 mL aliquot was thawed by adding 9 mL of PBS directly to the tube.

Confluency of 80% was targeted for optimal transfection. Cells were seeded either in 6-well or 12-well plates for transfection. Catalog numbers for 6, 12, 24, 48, and 96 well plates used at various points in these experiments are VWR CAT #10062-892, 10062-894, 10062-896, 10062-898, 10062-900. Cells were transfected using either Lipofectomene 3000 (ThermoFisher CAT #L3000008) or Polyplus JetOptimus (Polyplus-Sartorius CAT #101000006) according to manufacturer's protocols. DNA amounts were scaled according to the surface area of the well plate used. For example, the following amounts of DNA were used for a 6-well plate or a 35 mm diameter dish:

- Halo-TM-SNAP: 1 µg
- Halo-β2AR: 1 µg
- GαS-SNAP: 1 µg
- SNAP-CaaX: 2.5 µg
- Gβ: 0.5 µg
- Gγ: 0.2 µg

***A.3. Plasmids.*** The following plasmids were used. Full circular sequence maps for all plasmids are provided as downloadable files.

- pHR-CMV-Tet02_Twin-Strep: pHR-CMV-Tet02_Twin-Strep was purchased from Addgene (plasmid #113833).
- Halo-TM-SNAP was received as a gift from Dr. Johannes Broichhagen at Leibniz-Forchungsinstitut for Molecular Pharmocology (20). Amino acid sequence of Halo-TM-SNAP:

```
MVPSSDPLVTAASVLEFGLGISTMETDTLLLWVLLLWVPGSTGDYPYDVPDYAGAQPARSGSEIG
TGFPFDPHYVEVLGERMHYVDVGPRDGTPVLFLHGNPTSSYVWRNIIPHVAPTHRCIAPDLIGMG
KSDKPDLGYFFDDHVRFMDAFIEALGLEEVVLVIHDWGSALGFHWAKRNPERVKGIAFMEFIRPI
PTWDEWPEFARETFQAFRTTDVGRKLIIDQNVFIEGTLPMGVVRPLTEVEMDHYREPFLNPVDRE
PLWRFPNELPIAGEPANIVALVEEYMDWLHQSPVPKLLFWGTPGVLIPPAEAARLAKSLPNCKAV
DIGPGLNLLQEDNPDLIGSEIARWLSTLEISGVDEQKLISEEDLNAVGQDTQEVIVVPHSLPFKV
VVISAILALVVLTIISLIILIMLWQKKPRGAQPARSMDKDCEMKRTTLDSPLGKLELSGCEQGLH
EIIFLGKGTSAADAVEVPAPAAVLGGPEPLMQATAWLNAYFHQPEAIEEFPVPALHHPVFQQESF
TRQVLWKLLKVVKFGEVISYSHLAALAGNPAATAAVKTALSGNPVPILIPCHRVVQGDLDVGGYE
GGLAVKEWLLAHEGHRLGKPGLGGRLEVLFQGVD
```

- Halo-β2AR was purchased from Addgene (plasmid #66994) and transformed into backbone vector #113833. Amino acid sequence of Halo-β2AR:

```
MATGSRTSLLLAFGLLCLPWLQEGSAFPTIPLSGSEIGTGFPFDPHYVEVLGERMHYVDVGPRDG
TPVLFLHGNPTSSYLWRNIIPHVAPSHRCIAPDLIGMGKSDKPDLDYFFDDHVRYLDAFIEALGL
```

EEVVLVIHDWGSALGFHWAKRNPERVKGIACMEFIRPIPTWDEWPEFARETFQAFRTADVGRELI
IDQNAFIEGALPMGVVRPLTEVEMDHYREPFLKPVDREPLWRFPNELPIAGEPANIVALVEAYMN
WLHQSPVPKLLFWGTPGVLIPPAEAARLAESLPNCKTVDIGPGLFLLQEDNPDLIGSEIARWLPG
LAGSGGGGSMGQPGNGSAFLLAPNRSHAPDHDVTQQRDEVWVVGMGIVMSLIVLAIVFGNVLVIT
AIAKFERLQTVTNYFITSLACADLVMGLAVVPFGAAHILMKMWTFGNFWCEFWTSIDVLCVTASI
ETLCVIAVDRYFAITSPFKYQSLLTKNKARVIILMVWIVSGLTSFLPIQMHWYRATHQEAINCYA
NETCCDFFTNQAYAIASSIVSFYVPLVIMVFVYSRVFQEAKRQLQKIDKSEGRFHVQNLSQVEQD
GRTGHGLRRSSKFCLKEHKALKTLGIIMGTFTLCWLPFFIVNIVHVIQDNLIRKEVYILLNWIGY
VNSGFNPLIYCRSPDFRIAFQELLCLRRSSLKAYGNGYSSNGNTGEQSGYHVEQEKENKLLCEDL
PGTEDFVGHQGTVPSDNIDSQGRNCSTNDSLL

- GαS-SNAP was designed in our lab and synthesized by TWIST Biosciences. The gene was sub-cloned into backbone vector #113833 using Gibson Assembly. SNAP-tag placement was chosen according to functional studies with a BRET-based sensor (21). Amino acid sequence of GαS-SNAP:

MGCLGNSKTEDQRNEEKAQREANKKIEKQLQKDKQVYRATHRLLLLGAGESGKSTIVKQMRILHV
NGFNGEGGEEDPQAARSNSDGEKATKVQDIKNNLKEAIETIVAAMSNLVPPVELANPENQFRVDY
ILSVMNSGGGGTRSTGMDKDCEMKRTTLDSPLGKLELSGCEQGLHRIIFLGKGTSAADAVEVPAP
AAVLGGPEPLMQATAWLNAYFHQPEAIEEFPVPALHHPVFQQESFTRQVLWKLLKVVKFGEVISY
SHLAALAGNPAATAAVKTALSGNPVPILIPCHRVVQGDLDVGGYEGGLAVKEWLLAHEGHRLGKP
GLGGTATGGGGSVPDFDFPPEFYEHAKALWEDEGVRACYERSNEYQLIDCAQYFLDKIDVIKQAD
YVPSDQDLLRCRVLTSGIFETKFQVDKVNFHMFDVGGQRDERRKWIQCFNDVTAIIFVVASSSYN
MVIREDNQTNRLQEALNLFKSIWNNRWLRTISVILFLNKQDLLAEKVLAGKSKIEDYFPEFARYT
TPEDATPEPGEDPRVTRAKYFIRDEFLRISTASGDGRHYCYPHFTCAVDTENIRRVFNDCRDIIQ
RMHLRQYELL

- Gβ was purchased from Addgene (plasmid #133856). It was sub-cloned into backbone vector #113833 without the fluorescent tag. Amino acid sequence of Gβ:

MSELDQLRQEAEQLKNQIRDARKACADATLSQITNNIDPVGRIQMRTRRTLRGHLAKIYAMHWGT
DSRLLVSASQDGKLIIWDSYTTNKVHAIPLRSSWVMTCAYAPSGNYVACGGLDNICSIYNLKTRE
GNVRVSRELAGHTGYLSCCRFLDDNQIVTSSGDTTCALWDIETGQQTTTFTGHTGDVMSLSLAPD
TRLFVSGACDASAKLWDVREGMCRQTFTGHESDINAICFFPNGNAFATGSDDATCRLFDLRADQE
LMTYSHDNIICGITSVSFSKSGRLLLAGYDDFNCNVWDALKADRAGVLAGHDNRVSCLGVTDDGM
AVATGSWDSFLKIWN

- Gγ was purchased from Addgene (plasmid #133857). It was sub-cloned into backbone vector #113833 without the fluorescent tag. Amino acid sequence of Gγ:

MASNNTASIAQARKLVEQLKMEANIDRIKVSKAAADLMAYCEAHAKEDPLLTPVPASENPFREKK
FFCAIL

- SNAP-CaaX: pSNAP-CaaX was purchased from Addgene (plasmid #101133). Amino acid sequence of SNAP-CaaX:

MELHRGGGRDIKLTMDKDCEMKRTTLDSPLGKLELSGCEQGLHEIKLLGKGTSAADAVEVPAPAA
VLGGPEPLMQATAWLNAYFHQPEAIEEFPVPALHHPVFQQESFTRQVLWKLLKVVKFGEVISYQQ
LAALAGNPAATAAVKTALSGNPVPILIPCHRVVSSSGAVGGYEGGLAVKEWLLAHEGHRLGKPGL
GPAGCMSCKCVLS

#### B. Confocal imaging

***B.1. Fluorescent labeling of cells for confocal imaging.*** We used coated glass-bottom dishes directly from the manufacturer with no additional cleaning. A list of dishes used: Matek CAT #P35G-1.5-14-C, VWR CAT #75856-742, Cellvis CAT #d35-14-1.5-n. Cells were seeded onto glass-bottom dishes and labeled with Janelia Fluor (JF) dyes. The dyes used were JF479-HaloTag, JF552-HaloTag, JF552-cpSNAP-tag, JF525-HaloTag, JF549i-SNAP-tag, JF549i-HaloTag, and JFX646-SNAP-tag. Identities of dyes used in specific experiments can be found in figure captions. Long-term, all dye stocks were stored at 500 µM at -20°C, and at 4°C in a dark box for short-term storage.

For cell labeling, a dye solution in cell media (DMEM + 10% FBS + PS) was prepared at a final concentration of 2 µM. Solution volume was sufficient to fully cover the bottom surface of the dish (e.g., 500 µL for a 35 mm dish). For the labeling reaction, the dye solution was warmed to 37°C and cells were incubated with this solution at 37°C for 15-20 minutes, then washed. The protocol for one wash cycle was as follows:

- Dye/DMEM solution was aspirated.
- PBS at 37°C was added gently and swirled. Enough solution was added to cover the bottom of the dish completely.
- PBS was aspirated.
- Steps 2 and 3 were repeated once.
- Fresh cell media (DMEM + 10% FBS + PS) was added and cells were placed in the incubator.

After a 15-minute incubation, the entire wash cycle was repeated but in the final wash step, phenol-red-free FluoroBrite DMEM was used. Next, confocal imaging was performed.

We found that without an incubation step between wash cycles, background dye remained inside the cell. Additional incubation time allowed this non-specifically bound dye to diffuse out of the cell, which improved imaging contrast.

***B.2. Confocal image acquisition.*** Confocal image acquisition was performed at the Harvard Center for Biological Imaging (HCBI) using a Zeiss LSM 880 confocal microscope with a Zeiss Plan-Apochromat 63x / 1.40 NA oil DIC M27 objective (Zeiss CAT #420782-9900-000). Immersol 518F imaging oil (Microscope World CAT #444960-0000-000) was used. Images were acquired at a pixel depth of 16 bits. Images were acquired in frame scanning mode with line averaging set to 2 per pixel. Pixel dwell time was 12.20 µs for Halo-TM-SNAP with a pixel size of 98 nm, and 16.38 µs for Halo-β2AR/SNAP-CaaX with a pixel size of 88 nm. Green fluorescence was excited with a 488 nm multi-line argon laser. Red fluorescence was excited with a DPSS 561 nm laser. Far-red fluorescence was excited with a HeNe 633 nm laser. The lasers were directed to the objective via a 458/561 multi-beamsplitter for green and red excitation and a 514/561/633 multi beamsplitter for far red excitation. The following laser powers and gain settings were used to acquire images presented in figures:

Halo-β2AR/SNAP-CaaX:
561: laser power 0.056 mW (~44 kW/cm$^2$), gain 750, detection window 570-651 nm
633: laser power 0.062 mW (~39 kW/cm$^2$), gain 800, detection window 638-755 nm

Halo-TM-SNAP:
488: laser power 0.165 mW (~173 kW/cm$^2$), gain 811.4, detection window 490-570 nm
561: laser power 0.282 mW (~224 kW/cm$^2$), gain 650, detection window 566-679 nm

The pinhole was set to 1 AiryUnit. The spectral point detector used was a 32 channel GaAsP PMT for green and red emission plus the IR enhanced PMT 2 for far red emission.

***B.3. Confocal image processing.*** Confocal images were adjusted for brightness and contrast using Fiji software. All original images are available upon request.

#### C. SMLM imaging

***C.1. Preparation of mounting surfaces.*** We mounted coverslips on plain pre-cleaned glass slides, 25 x 75 mm, 1 mm thick (VWR CAT #48300-026). Glass slides were cleaned with methanol (Sigma Aldrich CAT #34860-1L-R) right before mounting to remove any dust. The mounting surface was covered with methanol using a dropper pipette and wiped once in one direction with a KimWipe. Any remaining methanol quickly evaporated, and the slides were ready for mounting. We used 22 x 22 mm No. 1.5 glass coverslips (VWR CAT #48366-227). Coverslips were placed on a Wash-N-Dry coverslip rack (Sigma Aldrich CAT #Z688568) to expose both sides of the surface. The rack was placed in a glass beaker and submerged in 1 M HCl. The beaker was covered with 2 layers of aluminum foil and heated to 60°C for 4-6 hours. Coverslips were then washed twice by immersing the entire coverslip rack into fresh DI water each time. Finally, coverslips were washed one additional time using 100% ethanol. The coverslips were stored in ethanol for up to 1 hour. Next, coverslips were removed from 100% ethanol and allowed to dry inside of a sterile TC hood. One coverslip was placed in each well of a 6 well tissue culture plate and 2 mL of DMEM + 10% FBS + PS were added.

***C.2. Fluorescent labeling.*** Cells seeded in a multi-well plate were labeled with photo-activatable (PA) Janelia Fluor (JF) dyes. We used PA-JF549-HaloTag and PA-JF646-SNAP-tag dyes. Identities of dyes used in specific experiments can be found in figure captions. All dyes were stored at 500 µM at -20°C for long-term storage. In contrast to confocal dyes, SMLM dyes were not stored at 4°C.

For cell labeling, a dye solution in cell media (DMEM + 10% FBS + PS) was prepared at a final concentration of 2 µM. Solution volume was sufficient to fully cover the bottom surface of the dish (e.g., 500 µL for a 35 mm dish). For the labeling reaction, the dye solution was warmed to 37°C and cells were incubated with this solution at 37°C for 15-20 minutes, then washed. The protocol for one wash cycle was as follows:

- Dye/DMEM solution was aspirated.
- PBS at 37°C was added gently and swirled. Enough solution was added to cover the bottom of the dish completely.
- PBS was aspirated.
- Steps 2 and 3 were repeated once.
- Fresh cell media (DMEM + 10% FBS + PS) was added and cells were placed in the incubator.

After a wash cycle cells were allowed to incubate at 37°C for 15 minutes. We repeated this wash cycle with the 15-minute incubation step 6 total times. Trypsin + EDTA (prepared as described in "B. Cell Culture and Transfection") was added to detach cells. Next, cells were seeded at low confluency (∼25%) onto the HCl-washed coverslips (from "C.1. Preparation of Mounting Surfaces").

***C.3. Fixation buffer.*** Fixation buffer was prepared using 4% w/v granular paraformaldehyde (PFA) (Electron Microscopy Sciences CAT #19208) in PBS (∼50-500 mL total volume). PFA/PBS mixture was heated to 50-60°C using a heating plate while stirring with a stir bar. Initially, the solution was cloudy as not all PFA dissolved. Next, 1 M NaOH in Milli-Q water was added drop-wise until the PFA/PBS solution turned clear. The solution was stirred at 50-60°C until all PFA dissolved, which typically took 5-10 minutes. The pH was adjusted to 7.4 and the solution was frozen at -20°C in 10 mL aliquots in 15 mL conical centrifuge tubes. When needed, aliquots were thawed in water at room temperature and sonicated until completely clear. Once thawed, aliquots were not refrozen.

***C.4. Fixation protocol.*** Media was aspirated from dishes with fluorescently labeled cells (from "C.2. Fluorescent Labeling") and replaced with fixation buffer. Cells were then kept in fixation buffer at room temperature for 15 minutes. During this process the dishes were swirled gently every 5 minutes. After 15 minutes of fixation, cells were washed with room-temperature PBS three times and stored at 4°C if not imaged immediately.

Immediately before imaging, PBS was gently aspirated. 0.1 µm TetraSpek Microspheres (Thermo-Fisher CAT #T7279) were added to the glass coverslip containing cells to act as fiducial markers. The concentration from the manufacturer was reported to be $\sim 1.8 \times 10^{11}$ particles/mL. We diluted Microspheres by a factor of 200 in Milli-Q water and sonicated before each use. This dilution factor led to the desired surface density of Microspheres and was determined empirically by serial dilution and imaging. We aimed for 5-10 Microspheres per field of view. 10 µL of diluted Microspheres were added to each coverslip. Diluted Microspheres were stored at 4°C.

***C.5. Mounting coverslips with cells for SMLM imaging.*** To mount coverslips, a few µL of Fluoromount-G mounting media (ThermoFisher CAT #00-4958-02) were added to the cleaned microscope slide (from "C.1. Preparation of Mounting Surfaces"). Fluoromount-G is extremely viscous. To ensure there were no bubbles, 20 µL of Fluoromount-G were drawn up in a pipette and a small portion was expelled. Next, a drop of Fluoromount-G was added to the microscope slide. A fresh pipette tip was then used to remove any air bubbles within the deposited drop of Fluoromount-G. Coverslips with cells facing the mounting media were mounted on top of Fluoromount-G and allowed to set for 10 minutes. After 10 minutes, the edges of the coverslips were sealed with clear nail polish and allowed to dry for ∼10 minutes. The samples were then moved to a light-proof slide storage box. Samples were stored at 4°C.

***C.6. SMLM image acquisition.*** SMLM image acquisition was performed on a Zeiss Elyra microscope at the HCBI with a Zeiss Plan-Apochromat 63x / 1.40 NA oil DIC M27 objective. Immersol 518F was used.

PA-JF549-HaloTag was excited with a DPSS 561 nm laser. 561 nm laser power measured after the objective without oil was 30.1 mW. With the objective removed, the power at the back focal plane was 40.3 mW. In parallel with 561 nm excitation, PA-JF646-SNAP-tag was excited with a 642 nm HeNe laser. HeNe laser power measured after the objective without oil was 30.2 mW. With the objective removed, the power at the back focal plane was 44.0 mW. Under these illumination conditions, the on-time of a single dye molecule was between 1 and 3 frames (100-300 ms). The sample was also illuminated with a 405 nm photo-activation laser between 561 nm and 641 nm excitation and collection. A schematic of a single frame of imaging is provided in Fig. S9. 405 nm laser power was increased over the duration of the experiment – as localizations became less frequent – from 0.031 mW to 0.626 mW measured after the objective without oil, corresponding to 0.052 mW and 1.04 mW respectively in the back focal plane.

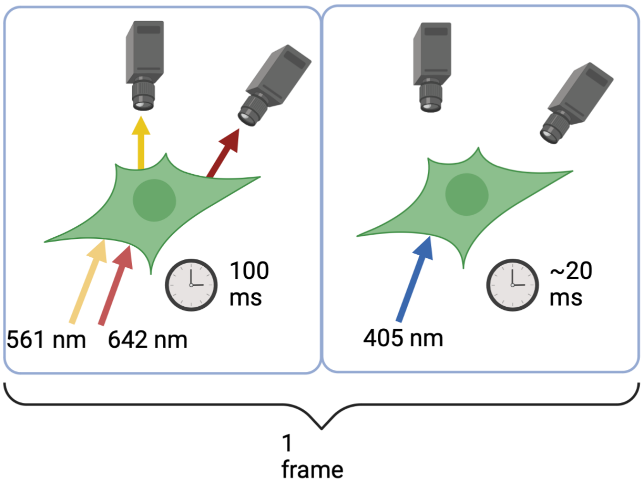

**Figure S9.** A single frame of SMLM illumination. Each frame has an exposure time of 100 ms for the 561 and 642 nm lasers. For approximately 20 ms during frame transfer and camera data processing, 405 nm laser light stochastically activates more fluorophores. 405 nm light is not present during camera acquisition.

The sample was initially illuminated with 561 nm and 642 nm laser lines (without 405 nm illumination) to photo-bleach prematurely photo-activated fluorophores. After 50-100 frames, the 405 laser was turned on.

Emission from both dyes was collected on two spatially registered cameras. The main beamsplitter for both channels was a quad 405/488/561/642 mirror. The emission was directed to two cameras through a DuoLink unit using a band-pass 490-560 + long-pass 640 dichroic camera splitter that reflected red emission (560-640) to camera 1 and far red emission (640+) to camera 2. The cameras used were Dual PCO.edge 4.2 CLHS sCMOS camera. The frame exposure time was set to 100 ms at 16 bits. The frame size was 49x49 µm.

### Supplementary Note 10: Analysis of experimental SMLM data

#### A. Preprocessing experimental SMLM images

***A.1. Generating localizations from SMLM images.*** SMLM images were processed using Zeiss Zen software to identify localizations. To select localizations, peak mask size was set to 9 pixels and fit a Gaussian mask over potential localizations. Peak intensity-to-noise ratio was set to 6, which selected among potential localizations and kept localizations if the standard deviation of its intensity against the rest of the image was above 6. Overlapping localizations (as defined by the mask of an event) were discarded. Localizations were filtered by localization precision (7 to 40 nm), number of photons (700-8000 photons) and experimental PSF size (90-180 nm). Localization precision was estimated by Zen using a Monte-Carlo Simulation of the experimental PSF including noise. These localization filters were selected based on respective localization histograms. They primarily excluded out-of-focus fluorophores and any dim fluorescent background. Spatially correlated localizations were grouped to aggregate emission of a single fluorophore that persisted over multiple camera frames. Localizations were then corrected for drift as described below.

***A.2. Drift correction.*** Typical SMLM data acquisition lasted from 20 to 40 minutes per image. To characterize drift, we imaged 100 nm TetraSpek Microspheres mounted between a glass slide and a coverslip as described above. We acquired 10,120 frames over approximately 33 minutes in two colors and observed drift on the order of several hundred nanometers to microns (Fig. S10(a)). Drift was corrected using Zen's model-based drift correction algorithm. Zoomed-in images of a bead before and after drift correction are shown in Fig. S10(b). After drift correction was applied, the full width at half maximum of the cluster of localizations was 18 nm for the green channel and 29 nm for the magenta channel.

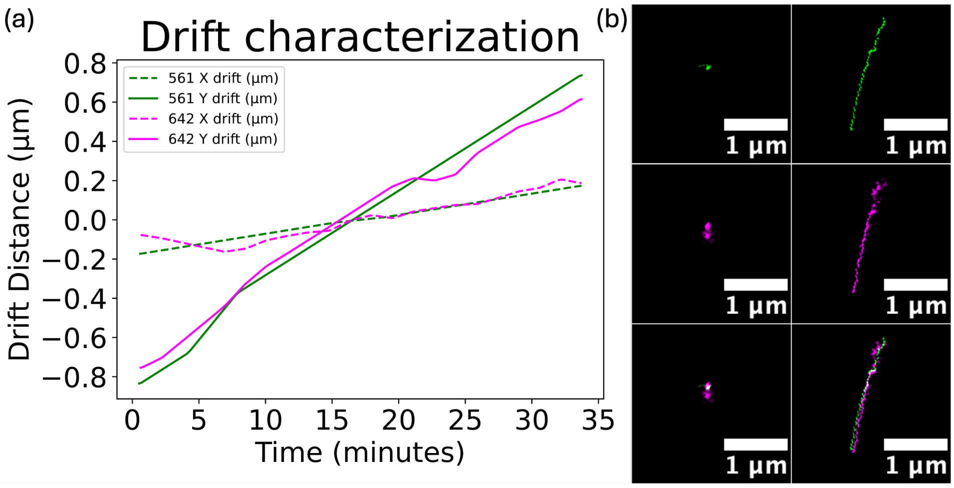

**Figure S10.** Model based drift correction algorithm was applied to the imaging signal derived from a 100 nm TetrakSpek fluorescent Microsphere. (a) Drift in the $x$ and $y$ direction for both 561 and 642 nm channels. (b) Drift correction as a function of frame number was applied in each channel. Before and after images of the two channels show the drift in the two channels as well as the efficacy of the correction.

***A.3. Removing areas of high density and dividing into subregions.*** Next, we generated a scatter plot based on the positions of drift-corrected localization signals in each cell. In order to remove extraneous localizations outside of the cell, we drew a boundary made of line segments to approximate the shape of the cell. The resulting polygon defined $\Omega$, the effective region of analysis. Any localizations outside of $\Omega$ were discarded from further analysis. Next, we eliminated localization signals that were part of tight clusters, using the clustering algorithm DBSCAN (22), with parameters eps $= 75$ nm and min_samples $= 10$.

Then we divided $\Omega$ into smaller subregions, by overlaying $\Omega$ with $d \times d$ equally sized boxes, where by default $d = 10$. See Fig. S11 for an example. Each subregion $R_{ij}$ was defined as where $\Omega$ and one of these boxes overlap, with effective area $S_{ij}$. Any subregion, where in at least one of two channels the density was $> 35\,\mu\text{m}^{-2}$, was again discarded.

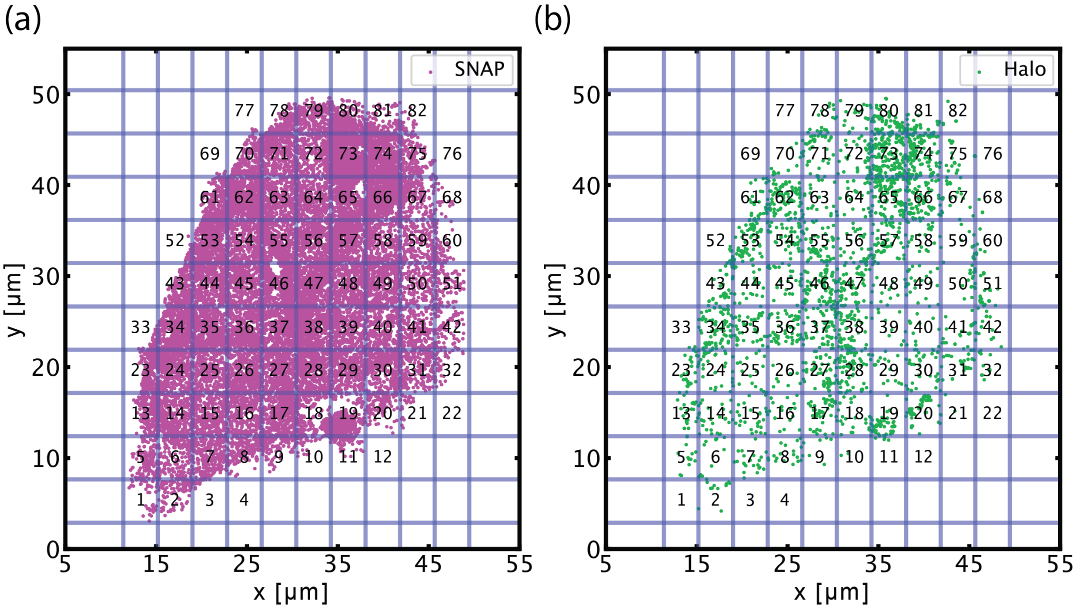

**Figure S11.** Divide into subregions by overlaying with $d \times d$ boxes ($d = 10$). (a) SNAP localizations, and (b) Halo localizations in a cell expressing Halo-TM-SNAP.

#### B. Analysis of experimental SMLM dataset

On each subregion $R_{ij}$, we applied our pipeline (Fig. S4) to the $N_{A,ij} + N_{B,ij}$ localizations to estimate the number of coupled pairs $N_{\text{coupled},ij}$ in $R_{ij}$. Note that in principle, iMEC can estimate more background pairs than the number of pairs in a subregion (i.e., $R_{ij}$ has less couplings than uniform random distributions of signals), in which case $N_{\text{coupled},ij} = 0$. We calculated the % of coupling as

$$q_{ij} = \frac{N_{\text{coupled},ij}}{\min(N_{A,ij}, N_{B,ij})}.$$

In Fig. 5 of the main text we reported the average $\langle q_{ij} \rangle$ for each cell.

The average over entire cells supported the same qualitative conclusion as from considering subregions with similar densities, that the percentage of couplings in cells expressing Halo-TM-SNAP was significantly higher than cells expressing Halo-β2AR/SNAP-CaaX. In Fig. S12, we plotted $\langle q_{ij} \rangle$, averaged over subregions with similar densities. More specifically, these were subregions whose densities $\min\{\rho_{A,ij}, \rho_{B,ij}\}$ lie in [0,4], [4,8], [8, 12], or [12, 16] µm$^{-2}$ respectively. The percentage $\langle q_{ij} \rangle$ displayed larger variability between different density ranges, but overall, the percentage of coupling is significantly higher for cells expressing Halo-TM-SNAP than those expressing Halo-β2AR/SNAP-CaaX.

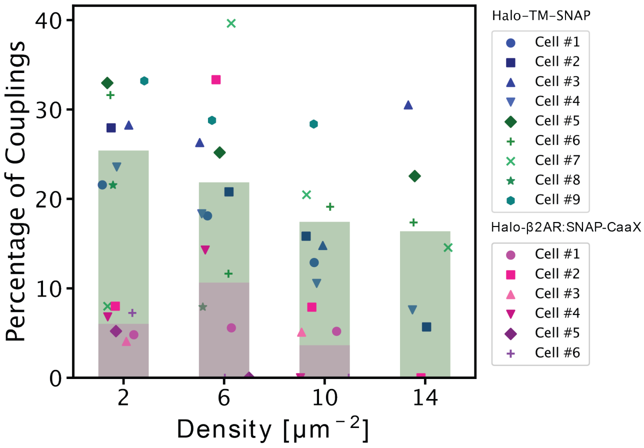

**Figure S12.** Percentage of couplings in subregions of certain densities (each marker represents a single cell). Across all densities, more couplings were identified in cells expressing Halo-TM-SNAP than those expressing Halo-β2AR/SNAP-CaaX.

#### C. Comments on choices made in the analysis

***On dividing into subregions.*** Different local regions of the same cell spanned a range of densities. We opted to run the analysis with $d = 10$ as well as $d = 1$. With $d = 10$ (Fig. 5 of the main text), the % of couplings for both cells lines were nearly identical to when $d = 1$ (Fig. S13), with 21% couplings identified for Halo-TM-SNAP and 6% for Halo-β2AR/SNAP-CaaX.

***On removing clusters and subregions with high densities.*** Clusters could have arisen for various reasons, including overcounting or biological factors. Biological factors include cellular recycling into vesicles and endosomal pockets that may trap non-specifically bound dye. These clusters exhibited significantly elevated local densities, typically exceeding 500 µm$^2$, which is beyond the range in which our algorithm performs well. Thus, clusters were removed by DBSCAN prior to analysis. Similarly, subregions with densities greater than $35\,\mu\text{m}^{-2}$ were excluded from analysis to maintain a certain level of performance.

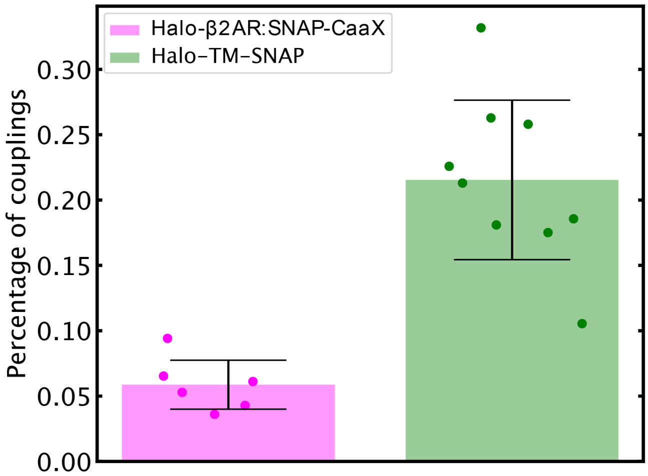

**Figure S13.** Percentage of couplings without dividing the cell into subregions (6 cells used for Halo-β2AR/SNAP-CaaX, and 9 cells used for Halo-TM-SNAP). Error bars indicate ±1 standard deviation.

### Supplementary Note 11: Comments on analysis of experimental images

In an ideal scenario, we would expect the percentage of couplings for Halo-TM-SNAP to be 100%, and 0% for Halo-β2AR/SNAP-CaaX. The experimental data deviated from this scenario, with only ∼20% of couplings identified for cells expressing Halo-TM-SNAP (Figs. 5(d), S12, S13), and in certain subregions, >0% for Halo-β2AR/SNAP-CaaX (Fig. S12). In this section we discuss potential explanations for these observations.

#### A. Halo-TM-SNAP

The protein Halo-TM-SNAP is a transmembrane protein recombinantly fused to a HaloTag and a SNAP-tag (Fig. 5(a, right) of main text). Thus, ideally, we should have observed 100% couplings, however, not all tags were observed. The missing localizations could be ascribed to several causes. First, the labeling efficiency and the detection rate were not perfect. Second, certain localizations were excluded in the preprocessing step, based on localization precision, point spread function half-width, and what were believed to be instances of overcounting. Third, some dye molecules were activated prior to imaging, which was started after ∼100 frames.

In a positive control such as Halo-TM-SNAP, missing localizations affect the percentage of couplings identified. Consider for example, if only the HaloTag failed to be identified in a protein, the SNAP-tag that was detected might be paired with the HaloTag of a different protein by our algorithm. This situation may become too complex to parse in terms of the weighted bipartite graph in GMO. In Section C below, we showed that if molecules are detected at rate $1-r$ and the true number of couplings is $N$, then the expected number of couplings identified by our algorithm would be $N(1-r)^2$. For example, a detection rate of $r = 55\%$ would lead to a percentage of couplings ∼20%.

#### B. Halo-β2AR/SNAP-CaaX

Halo-β2AR and SNAP-CaaX are not expected to interact. Our pipeline identified a positive percentage of couplings for certain subregions of some cells, even though the expected percentage of couplings is 0%. In these subregions, we noticed that the two proteins were not uniformly distributed in space, but there were clusters. However, these clusters were not eliminated by DBSCAN (Fig. S14), which we suspect was due to our choice of parameters (eps=75 nm and min_samples=10). We also observed that when these clusters were present, localizations in both spectral channel were also dense in the same subregion, as demonstrated by Fig. S14.

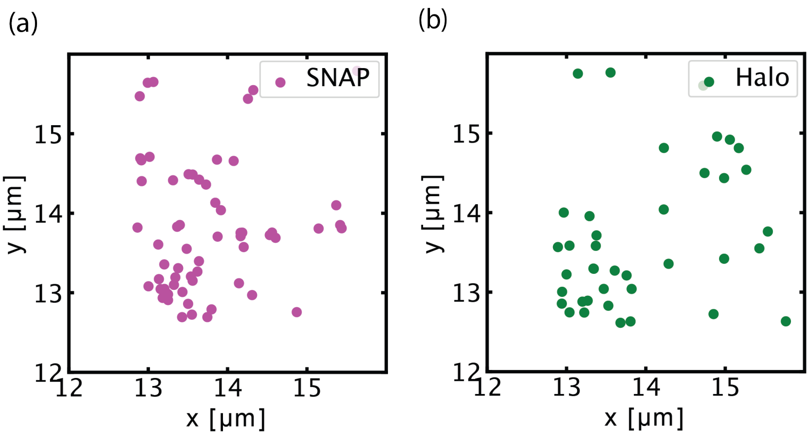

**Figure S14.** A subregion in a cell expressing Halo-β2AR/SNAP-CaaX. DBSCAN was not able to eliminate some clusters. The two channels also appeared to be correlated in terms of density in this subregion.

##### C. The estimated number of couplings scales with the square of detection rate

Here we considered the effect of missing detection on the percentage of couplings. In a system with $N_{\text{coupled}}$ couplings, if localizations of $A$ and $B$ were dropped at rate $r$, i.e., a detection rate of $1-r$, then in theory, the detected number of couplings is expected to be $N_{\text{coupled}}(1-r)^2$, which was the conclusion from our numerical experiments.

We conducted two numerical experiments at varying densities. In the first, we modelled a positive control system like Halo-TM-SNAP ($N_A = N_B = N_{\text{coupled}}$); in the second, we assumed equilibrium between the interacting partner and the complex ($N_A = N_B = 2N_{\text{coupled}}$). In each experiment, we simulated SMLM datasets, then randomly deleted localizations of $A$ and $B$ at rate $r$, and ran our algorithm on the resulting datasets. The parameters used can be found in Table S7. Fig. S15 shows that in both numerical experiments, the numbers of couplings detected by our pipeline matched the theoretical prediction $N_{\text{coupled}}(1-r)^2$ closely. Note that in these numerical experiments, $N_A$ and $N_B$ were kept constant, so the number of couplings was proportional to the percentage of couplings.

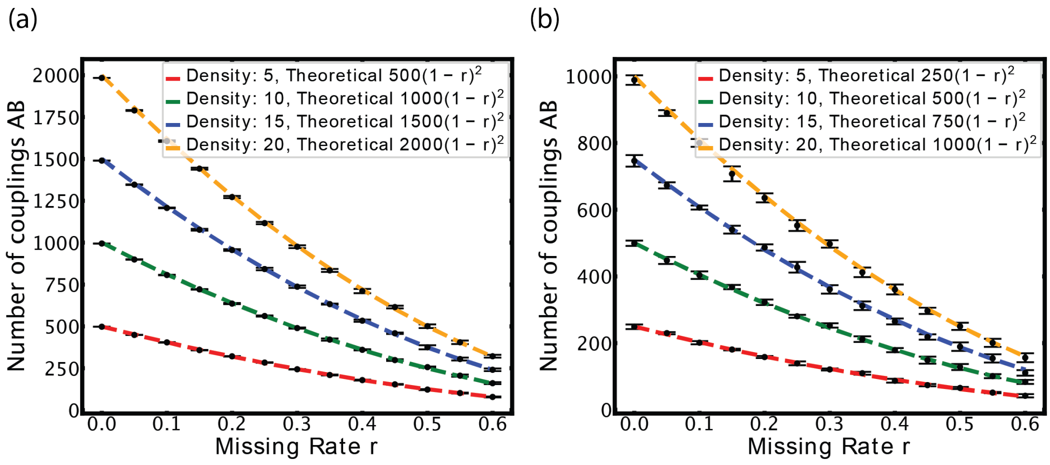

**Figure S15.** The number of couplings identified scales as $(1-r)^2$ where $1-r$ is the detection rate. (a) Positive control, where all molecules are pairs. (b) Equilibrium, where half the population are paired. Dashed Line: Theoretical Expression; Dots: Algorithm-Predicted Values. Error bars indicate ±3 standard deviations across 10 trials of iMEC.

**Table S7.** Parameters used to generate Fig. S15

| | |
|---|---|
| Density $\rho_A = \rho_B = \rho$ | See Fig. S15 |
| Area $S$ | $S = 100$ µm$^2$ |
| Number of AB | $\rho S$ or $\rho S/2$ |
| Number of background A | 0 or $\rho S/2$ |
| Number of background B | 0 or $\rho S/2$ |
| True distance $d_{\text{true}}$ | Sampled from Unif(0, 10 nm) |
| Localization Precision | Sampled from Fig. S6 |
| $d_{\text{lower}}$ | 0 nm |
| $d_{\text{upper}}$ | 20 nm |
| Search distance factor threshold $C$ | 4 |
| Number of sampling points to estimate probability | $10^5$ |
| Number of Monte Carlo steps to estimate $N^*_{\text{bg}}$ | 15 |
| Number of Monte Carlo trails in each estimation of $N^i_{\text{bg}}$ | 10 |

### Supplementary Note 12: Validation at chemical equilibrium

To see whether the algorithm could be used to infer the extent of couplings at the equilibrium, we applied it to the system $A + B \rightleftharpoons AB$ at equilibrium with equilibrium constant $K_{\text{eq}}$. The percentage of couplings is proportional to the concentration of $AB$, whose analytical expression can be derived:

$$[AB] := \frac{N_{AB}}{S} = \rho + \frac{1}{2K_{\text{eq}}} - \sqrt{\frac{1}{4K_{\text{eq}}^2} + \frac{\rho}{K_{\text{eq}}}}, \quad \textbf{(S6)}$$

where the density $\rho$ refers to the total amount over an area $S$, i.e., $\rho = (N_A + N_{AB})/S$ and $\rho = (N_B + N_{AB})/S$. We generated simulated dataset for $K_{\text{eq}} = 0.1, 0.5, 1$ and $\rho$ up to $100$ molecules µm$^{-2}$. All parameters used can be found in Table S8.

As shown in Fig. S16, error remained below 3.5% across all densities and equilibrium constants. The error exhibited a slight tendency to grow as density increased. Note that when the number of interacting partners $N_{\text{AB}}$ was small (e.g., for $K_{\text{eq}} = 0.1$ and low density), the large error was due to a small denominator in the definition of error.

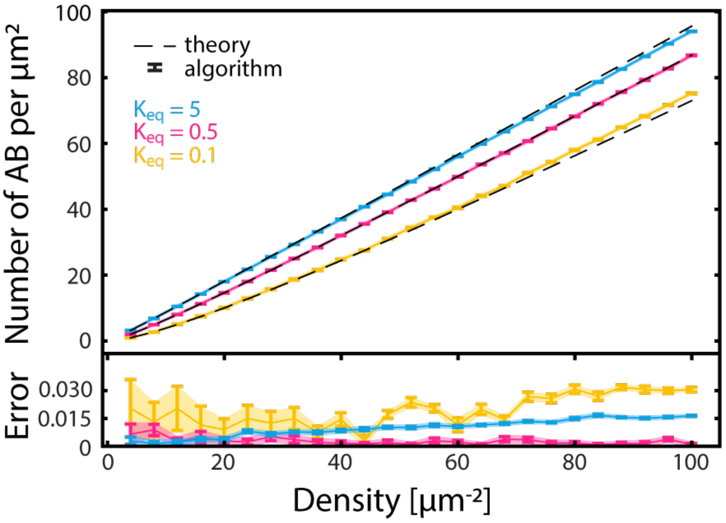

**Figure S16.** (Top) The density of $AB$ estimated by the algorithm (markers) and the theoretical density (dashed lines) at different $K_{\text{eq}}$ and densities. (Bottom) Error rate. Error bars indicate ±1 standard deviation across 10 trials of iMEC.

**Table S8.** Parameters used to generate Fig. S16

| | |
|---|---|
| Density $\rho$ | Varying $\rho$ (see $x$-axis) |
| Area $S$ | $S = 100\ \mu\text{m}^2$ |
| Number of AB | $N_{AB}$ (from Eq. S6) |
| Number of background A | $\rho S - N_{AB}$ from |
| Number of background B | $\rho S - N_{AB}$ from ) |
| True distance $d_{\text{true}}$ | Sampled from Unif(0, 10 nm) |
| Localization Precision $\sigma_A$, $\sigma_B$ | Sampled from Fig. S6 |
| $d_{\text{lower}}$ | 0 nm |
| $d_{\text{upper}}$ | 20 nm |
| Search distance factor threshold $C$ from SN 3 | 4 |
| Number of sampling points to estimate probability $N$ from Algorithm 1 | $10^5$ |
| Number of iterations $T$ to estimate $N^*_{\text{bg}}$ from Algorithm 2 | 25 |
| Number of MC trials $M$ in each estimation of $N^i_{\text{bg}}$ in Algorithm 2 | 10 |

### Supplementary Note 13: The algorithm can estimate the level of coupling accurately across different density ratios

Colocalization analysis has been notably affected by density fluctuations. Measuring the degree of coupling becomes challenging when there are variations in the density ratio between $A$ and $B$ (23; 24; 15). A comprehensive assessment requires comparisons across diverse cells with varying densities of molecules $A$ and $B$. Imagine a scenario where molecules of $A$ (with density $\rho_A$) colocalize with molecules of $B$ (with density $\rho_B$) at some percentage of colocalization. Now, if we increase the densities to $2\rho_B$ and $3\rho_B$ while maintaining the same percentage of couplings, an ideal colocalization algorithm should consistently yield the same level of couplings for these different density ratios. However, achieving this ideal outcome has proven challenging. Several approaches on incorporating densities to estimate the level of colocalization (using different definitions of colocalization) have been recently developed (14; 15).

Recall that we defined the percentage of couplings $q$ as the number of coupling molecules divided by the minimum of the number of molecules of $A$ and $B$. Here we would like to simulate a percentage of couplings $\gamma$ to either be $3\%$ (low binding) or $20\%$ (high binding), and see whether our pipeline returns a percentage of coupling $\langle q \rangle$ that is close to $\gamma$ or not. In a region with area $S$, $N_A$ and $N_B$ molecules of $A$ and $B$ respectively, i.e., at densities $\rho_A = N_A/S$, $\rho_B = N_B/S$, one would expect the number of coupling to be $N_{AB} = S\gamma \min\{\rho_A, \rho_B\}$. Correspondingly, the number of uncoupled $A$ would be $N_A = (\rho_A - \gamma \min\{\rho_A, \rho_B\})S$. A similar calculation for the number of coupled $B$ gives $N_B = (\rho_B - \gamma \min\{\rho_A, \rho_B\})S$. Thus, for different combinations of $\rho_A$ and $\rho_B$, we generated a dataset with $N_A$, $N_B$, and $N_{AB}$ molecules of $A$, $B$ and $AB$ respectively over a box of area $S$. Then we applied our algorithm and calculated the percentage of coupling $q(\rho_A, \rho_B)$. All parameters used are shown in Table S9.

A good algorithm would return a percentage of coupling $q(\rho_A, \rho_B)$ that is independent of $\rho_A, \rho_B$. Indeed, this is what we found; in Fig. S17 we plotted $q(\rho_A, \rho_B)$ for different combinations of $(\rho_A, \rho_B)$, and found that $q(\rho_A, \rho_B)$ is approximately $\gamma$, which was set at either 3% or 20%.

**Table S9.** Simulation Parameters in Fig. S17

| | |
|---|---|
| Density $\rho_A$, $\rho_B$ | Varying (see $x$-axis) |
| Area S | $100\ \mu\text{m}^2$ |
| Percentage $\gamma$ | $\gamma = 3\%$ or $20\%$ |
| Number of AB | $S\gamma \min(\rho_A, \rho_B)$ |
| Number of background A | $(\rho_A - \gamma \min(\rho_A, \rho_B))S$ |
| Number of background B | $(\rho_B - \gamma \min(\rho_A, \rho_B))S$ |
| True distance | Sampled from Unif(0,10 nm) |
| Localization Precision | Sampled from Fig. S6 |
| $d_{\text{lower}}$ | 0 nm |
| $d_{\text{upper}}$ | 20 nm |
| Search distance factor threshold $C$ | 4 |
| Number of sampling points to estimate probability | $10^5$ |
| Number of Monte Carlo steps to estimate $N^*_{\text{bg}}$ | 15 |
| Number of Monte Carlo trails in each estimation of $N^i_{\text{bg}}$ | 10 |

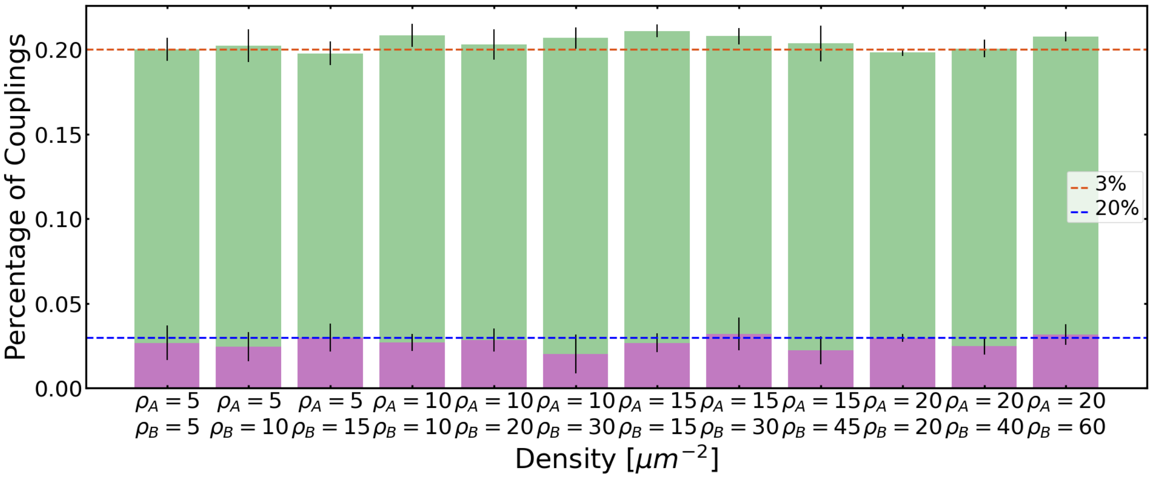

**Figure S17.** Estimated percentage of couplings $q$ under different combinations of densities for $A$ ($\rho_A$) and $B$ ($\rho_B$), where the dataset generated were consistent with percentage of couplings $\gamma = 3\%$ and 20% respectively. Error bars represent one standard deviation across 10 trials of iMEC.

### Supplementary Note 14: Inferring rate constants from kinetics data

We ask whether the algorithm can be used to infer kinetic parameters. We used the simulated non-equilibrium dataset in Fig. 6(d) of the main text, and the number of coupled molecules $N_{\text{coupled}}$ from our pipeline (GMO+iMEC) to estimate the rate constants $k_{\text{on}}$, $k_{\text{off}}$ of the binding reactions $A + B \rightleftharpoons AB$.

#### A. Analytical solution

In order to infer rate constants from our pipeline, we fitted the number of complexes $AB$ to the solution of the differential equations modelling this system. Consider the reaction $A + B \rightleftharpoons AB$ with rate constants $k_{\text{on}}$ and $k_{\text{off}}$ for the forward and backward reactions respectively. Under mass-action kinetics, with no $AB$ initially, the concentrations evolve according to the system of differential equations

$$\frac{d[A]}{dt} = \frac{d[B]}{dt} = -\frac{d[AB]}{dt} = -k_1[A][B] + k_2[AB],$$

with initial state $[A](0) = A_0$, $[B](0) = B_0$ and $[AB](0) = 0$. The solution is given by

$$\begin{aligned}[AB](t) &= \frac{A_0 B_0 \left(e^{-2\sqrt{c_2}k_1 t} - 1\right)}{(c_1 - \sqrt{c_2})e^{-2\sqrt{c_2}k_1 t} - c_1 + \sqrt{c_2}}, \\ [A](t) &= A_0 - [AB](t), \\ [B](t) &= B_0 - [AB](t),\end{aligned} \tag{S7}$$

where $c_1 = (A_0 + B_0 + k_2/k_1)/2$ and $c_2 = c_1^2 - A_0 B_0$.

#### B. Inferring rate constants

The rate constants $k_{\text{on}}$ and $k_{\text{off}}$ were obtained by fitting the density of $AB$ identified ($N_{\text{coupled}}/S$) at time $t$, where $S$ is the area of the simulation box, to $[AB](t)$ from Eq. (S7). Simulation and analysis parameters used can be found in Tables S1 and S5.

Compared to the true parameters $k_{\text{on}} = 0.0549$ molecules$^{-1}$µm$^2$s$^{-1}$ (see Fig. S5 and Supplementary Note 6B) and $k_{\text{off}} = 1$ s$^{-1}$, the rate constants from fitting were found to be

$$\begin{aligned}\hat{k}_{\text{on}} &= \frac{0.0535 + 0.0492 + 0.0489}{3} = 0.0505 \text{ molecules}^{-1}\mu\text{m}^2\text{s}^{-1}, \\ \hat{k}_{\text{off}} &= \frac{0.947 + 0.808 + 0.831}{3} = 0.862 \text{ s}^{-1},\end{aligned}$$

where we averaged over the three densities (“cases”) in Fig. 6(d) of the main text. Fig. S18 shows the fitted rate constants for each case separately. Note that estimation of the rate constants was more accurate at low density (Case I); this is consistent with our observation that the performance of our algorithm degrades with increasing density (Fig. 4(b) of the main text).

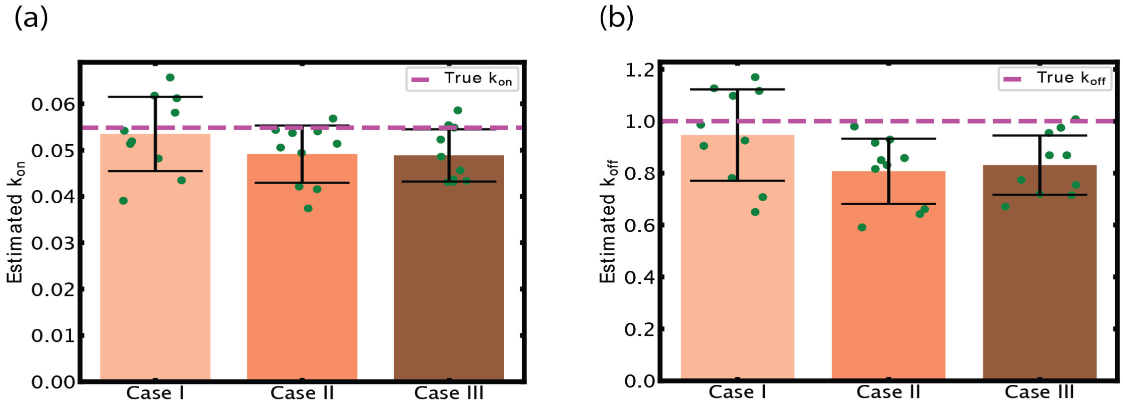

**Figure S18.** Inference of rate constants $k_{\text{on}}$ and $k_{\text{off}}$ from stochastic simulation, by fitting to Eq. (S7). (a) Estimated $k_{\text{on}}$ and (b) $k_{\text{off}}$ for Case I: $d_A = 10, d_B = 20$ µm$^{-2}$, Case II: $d_A = 20, d_B = 20$ µm$^{-2}$ and Case III: $d_A = 30, d_B = 20$ µm$^{-2}$. Error bars indicate $\pm 1$ standard deviation over 10 different trials.
